# Rejuvenation of aged oocyte through exposure to young follicular microenvironment

**DOI:** 10.1101/2024.03.11.584343

**Authors:** HaiYang Wang, Zhongwei Huang, Yaelim Lee, XinJie Song, Xingyu Shen, Chang Shu, Lik Hang Wu, Leroy Sivappiragasam Pakkiri, Poh Leong Lim, Xi Zhang, Chester Lee Drum, Jin Zhu, Rong Li

## Abstract

Reproductive aging is a major cause of fertility decline, attributed to decreased oocyte quantity and competence. Follicular somatic cells play crucial roles in the growth and development of the oocyte by providing nutrients and regulatory factors. Here we investigated how oocyte quality is affected by its somatic cell environment by creating chimeric follicles, whereby an oocyte from one follicle was transplanted into and cultured within another follicle whose native oocyte was removed. Somatic cells within the chimeric follicle re-establish connections with the oocyte and support oocyte growth and maturation in a three-dimensional (3D) culture system. We show that young oocytes transplanted into aged follicles exhibited reduced meiotic maturation and developmental potential, whereas the young follicular environment significantly improved the rates of maturation, blastocyst formation and live birth of aged oocytes. Aged oocytes cultured within young follicles exhibited enhanced interaction with somatic cells, more youth-like transcriptome, remodelled metabolome, improved mitochondrial function, and enhanced fidelity of meiotic chromosome segregation. These findings provide the basis for a future follicular somatic cell-based therapy to treat age-associated female infertility.

---

Female reproductive aging is associated with a sharp decline in the quantity and quality of eggs^1–3^. Post age 35, women exhibit elevated risks of miscarriage or childbirth affected by developmental disorders, primarily due to aneuploid eggs arising from chromosomal missegregation during oocyte maturation^4^. Unlike many somatic tissues that have self-renewing capacity, oocytes are highly specialized cells of which there is a finite pool deposited in the ovary during fetal development^5,6^. In many mammalian species, oocytes stay dormant for months to years before entering growth and maturation prior to fertilization. This developmental pattern is thought to be a cause of age-associated oocyte decline^3,6,7^. Despite decreasing fertility rates in many developed countries, there are no clinically viable methods to improve the developmental competence of aged oocytes ^2^, defined as the capacity to succeed in meiotic maturation, fertilization, and subsequent embryonic development. Current strategies, such as *in vitro* activation ^8,9^, intraovarian injection of platelet-rich plasma^10^, oocyte cryopreservation^11^, and ovarian tissue transplantation^12^, primarily address poor ovarian reserve and the decline in egg quantity. Although some studies suggest that mitochondrial replacement therapy may improve oocyte quality, its use is limited to a few countries due to ethical concerns, legal and regulatory challenges, and a lack of comprehensive understanding regarding its long-term safety and efficacy^13,14^.

The developmental competence of an oocyte is acquired in the ovarian follicle, the basic functional unit in the mammalian ovary composed of an oocyte surrounded by somatic cells that include granulosa cells (GCs) and theca cells^15,16^. The growth and maturation of ovarian follicles encompass several developmental stages, including primordial, primary, secondary, and antral follicles, culminating in the preovulatory stage. The secondary and antral stages are of particular importance as they involve noteworthy structural and cellular transformations within the follicle, essential for responding to hormonal signals required for growth and maturation^17–19^. During these stages, GCs provide many essential nutrients and metabolic precursors to the oocyte via transzonal projections (TZPs), actin-based extensions that connect GCs with the oocyte through gap junctions^16,20,21^. In turn, the oocyte provides signalling molecules important for the morphogenesis and proliferation of GCs. Through such exchange of regulatory and metabolic factors, the oocyte acquires developmental competence and is poised for successful fertilization after meiotic maturation^16,21,22^.

## Aged Follicular Environment Impairs Oocyte Quality

Although it is known that oocyte and GCs have a co-dependent relationship^16,23^, it is unclear how the somatic cell environments in follicles contribute to oocyte aging. To investigate this, we isolated follicles from young (2-3 month) and aged (14-15 month) mice to detect the aging induced dysfunction of surrounding somatic cells of oocytes, especially GCs. We found that the percentage of Ki-67-positive somatic cells, which reflects proliferative activity, was significantly lower in aged follicles compared to young follicles (Fig. 1a,b). Somatic cells in aged follicles also had higher levels of γH2AX and active caspase 3/7-positive cells, respectively as indicators of DNA damage and apoptosis, both being hallmarks of aging (Fig. 1c,d, Extended Data Fig. 1a,b). Additionally, aged follicular somatic cells exhibited reduced mitochondrial membrane potential (ΔΨm) and increased levels of reactive oxygen species (ROS) compared to young follicular somatic cells (Extended Data Fig. 1c-h). Interestingly, we observed a positive correlation of mitochondrial ΔΨm or ROS levels between follicular somatic cells and oocytes (Extended Data Fig. 1d,g). These findings suggest that follicular somatic cells accumulate age-related abnormalities.

**Figure 1.**
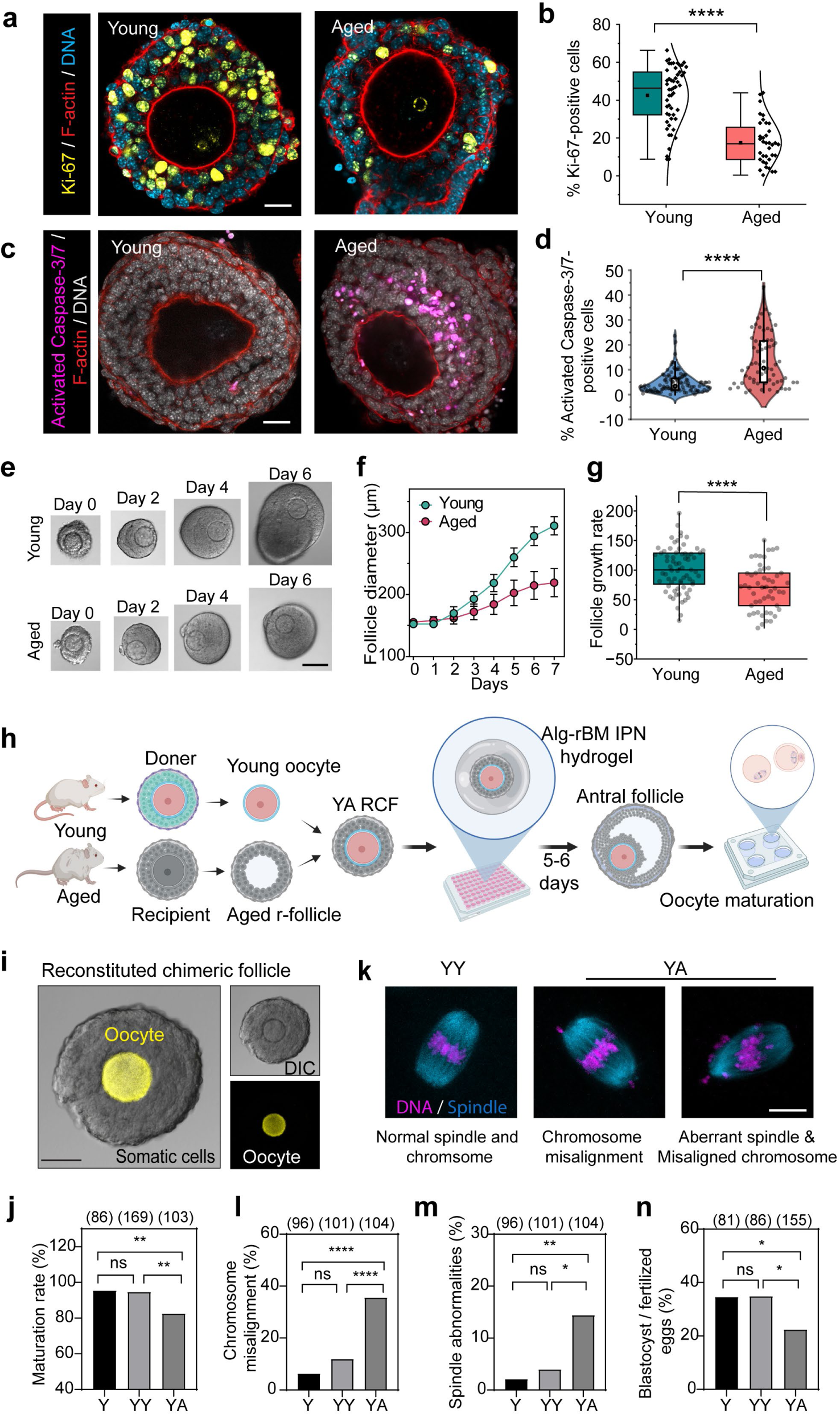
The impact of aged follicular environment on oocyte quality. **a.** Representative images of Ki-67 immunofluorescence in young and aged follicles. DNA was stained using DAPI, and F-actin was stained with phalloidin. Scale bar, 20 μm. **b.** Quantification of the percentage of Ki-67-positive cells per follicle. n = 53 (young), 36 (aged). A normal distribution is shown to the right of each box plot. Young follicles: 2-3 month-old females; aged follicles: 14-15 month-old females. **c.** Representative images of activated caspase-3/7 staining in young and aged follicles. DNA was stained using DAPI, and F-actin was stained with phalloidin. Scale bar, 20 μm. **d.** Quantification of the percentage of activated caspase-3/7-positive cells per follicle in young and aged follicles. n = 114 (young), 64 (aged). Young follicles: 2-3 month-old females; aged follicles: 14-15 month-old females. **e.** Sequential images of follicles from young (2 month) and aged (14 month) mice cultured in Alg-rBM IPN hydrogels. Scale bar, 100 μm. **f.** Diameters of follicles from young (2 months) and aged (14 months) mice monitored daily from day 0 to day 7. n = 18 (young), 18 (aged). For individual follicle tracking, please refer to Extended Data Fig. 2i. Data are shown as mean ± SEM. **g.** Box plots showing follicle growth rate in young (2-3 months) and aged (14-17 months) mice, calculated as (Follicle diameter at Day 7 - Follicle diameter at Day 0) / Follicle diameter at Day 0. n= 69 (young), 51 (aged). **h.** Schematic illustration of the RCF system using young oocyte transplanting to aged r-follicle as an example. **i.** Representative image of an RCF. To differentiate between the donor oocyte and r-follicle, we used a donor oocyte (pseudo-colored yellow) from transgenic mTmG mice with membrane-targeted tdTomato and an r-follicle (non-fluorescent) from a wild-type mouse. Only Fig. 1i involved mTmG mice; all other quantifications in Fig. 1 used only wild-type ICR mice. Scale bars, 50 μm. **j.** Comparison of oocyte maturation rates (PB1 extrusion rates) among oocytes from YY and YA RCFs as well as from young intact follicles (Y). The number of oocytes with the PB1 extruded was counted after maturation for 16h. ICR wide type mice were used. The numbers of oocytes are specified in brackets. Young oocytes: 2-month-old females; aged r-follicles: 14-month-old females. Fisher’s exact test. **k-m**. Aged r-follicles alleviated spindle and chromosomal alignment defects in young oocytes. (**k**) Representative live cell images of meiotic spindle and chromosomes in MII oocytes from YY and YA RCFs. The left image shows an example of normal spindle in young oocyte from YY RCF. Middle image shows an example of chromosomal misalignment in an oocyte from YA RCF, the image on right shows an example of abnormal spindle with chromosomal misalignment in an oocyte from YA RCF. Spindles (cyan) were stained with SiR-tubulin, and chromosomes (DNA, magenta) were stained with Hoechst. Scale bar, 10 µm. Quantification of the percentage of chromosomal misalignment (**l**) on the metaphase plate and spindle abnormalities (**m**) in oocytes from YY and AY RCFs as well as from young intact follicles (Y). The numbers of oocytes are specified in brackets. Young oocytes: 2-month-old females; aged r-follicles: 14-month-old females. Fisher’s exact test. **n.** Analysis of blastocyst formation rate following *in vitro* fertilization (IVF) of oocytes from YY and YA RCFs as well as from young intact follicles (Y). n= 81 (Y), 86 (YY), 155 (YA). Fisher’s exact test. Young oocytes: 2-month-old females; aged r-follicles: 14-month-old females. Box plots indicate mean and median by black square and center bar; upper and lower limits and whiskers represent quartiles and 1.5 × interquartile. Two-tailed unpaired t-tests for (**b**), (**d**), and (**g**). The data in (**j**), (**l**), (**m**), and (**n**) were analyzed by two-tailed Fisher’s exact test. \*\*\*\**P* < 0.0001; \*\**P* < 0.01; \**P* <0.05; ns, *P* > 0.05.

In order to study how the ageing of follicular environment affects oocyte quality and competence, we next established a 3D *ex vivo* culture system for follicle growth and development, building upon previous methods^24–27^. As illustrated in Extended Data Fig. 2a, secondary follicles from mouse ovaries were cultured and encapsulated in hydrogels of alginate-reconstituted basement membrane (rBM) interpenetrating network (Alg-rBM IPN), which maintained follicle morphology and enabled the follicles to increase in diameter and reach the antral (mature) stage (Extended Data Fig. 2b). We confirmed that oocytes grown in the follicle culture exhibited size and maturation rate (the frequency of first polar body (PB1) extrusion) and the ability to develop into blastocysts after fertilization comparable to those growing *in vivo* (Extended Data Fig. 2c-e). Additionally, the transcriptomes of *in vivo* and *in vitro* grown oocytes were similar (Extended Data Fig. 2f-h, Supplementary Table 1). These data suggest that our 3D *ex vivo* follicle culture system largely recapitulates the *in vivo* condition to support oocyte growth and development. We then used this culture system to investigate phenotypes associated with follicle ageing. Compared to follicles from young mice (2-3 months), follicles from aged mice (14-17 months) grew slower and showed higher levels of atresia in the 3D culture system (Fig. 1e-g, Extended Data Fig. 2i-k). These observations are consistent with the observed ageing phenotypes of the *in vivo* grown follicles.

To test whether the follicular somatic cell aging compromises the quality and developmental competence of oocytes, we established an experimental protocol whereby a denuded oocyte from a secondary follicle was transplanted into another secondary follicle whose own oocyte had been removed (referred to as recipient follicle, or r-follicle). The resulting structure is termed as reconstituted <u>c</u>himeric follicle (RCF) (Fig. 1h, Extended Data Fig. 3, Supplementary Video 1). Figure 1i shows an illustrative case where a doner oocyte from a secondary follicle of an mTmG mouse, which expresses a plasma membrane-targeted tdTomato, was transplanted into a r-follicle from a wild-type mouse; all other studies were conducted using wild-type mice unless specified otherwise. RCFs developed normally in our 3D culture system from secondary to the antral stage, and the oocytes within these RCFs matured similar to *in vitro* cultured intact follicles (Extended Data Fig. 4).

To assess whether follicular somatic cell aging affects oocyte quality, we transplanted <u>y</u>oung oocytes into <u>a</u>ged r-follicles (the resulting RCFs are designated YA) or <u>y</u>oung r-follicles (YY) as a control. Culturing in aged r-follicles significantly reduced the meiotic maturation of young oocytes (Fig. 1j). 3D confocal imaging revealed increased incidence of aberrant meiotic spindle and/or misaligned chromosomes in oocytes from YA RCFs compared to those from YY RCFs or unperturbed (not subjected to RCF manipulation) young follicles (Y) (Fig. 1k-m). Accompanying the reduced maturation capability and oocyte quality, YA RCFs-derived oocytes manifested elevated ROS and declined mitochondrial ΔΨm compared to those from Y or from YY RCFs (Extended Data Fig. 5). Furthermore, after *in vitro* fertilization (IVF), oocytes from YA RCFs exhibited reduced frequency of blastocyst formation compared to those from Y or YY RCFs (Fig.1n), indicating an impaired developmental potential of those young oocytes exposed to an aged follicular environment. As a side note, no significant differences were observed in the above measurements between oocytes from YY RCFs and Y (Fig. 1j-n, Extended Data Fig. 5b,d), suggesting that the RCF procedure itself did not impair oocyte quality.

## Improvement of Maturation and Developmental Competence of Aged Oocytes by Young Follicular Environment

We next investigated whether exposure to the young follicular environment could reduce defects of aged oocytes by transplanting them into <u>y</u>oung r-follicles (AY) or, as a direct control, <u>a</u>ged r-follicles (AA) following a similar protocol as the above (Fig. 2a). Culturing in young r-follicles significantly reduced the death rate (Extended Data Fig. 6a) and restored meiotic maturation of aged oocytes to a level similar to young oocytes YY RCFs (Fig. 2b). As assessed by confocal imaging, the occurrence of aberrant spindle and chromosome misalignment were significantly reduced in oocytes from AY, compared to AA, RCFs (Extended Data Fig. 6b-d).

**Figure 2.**
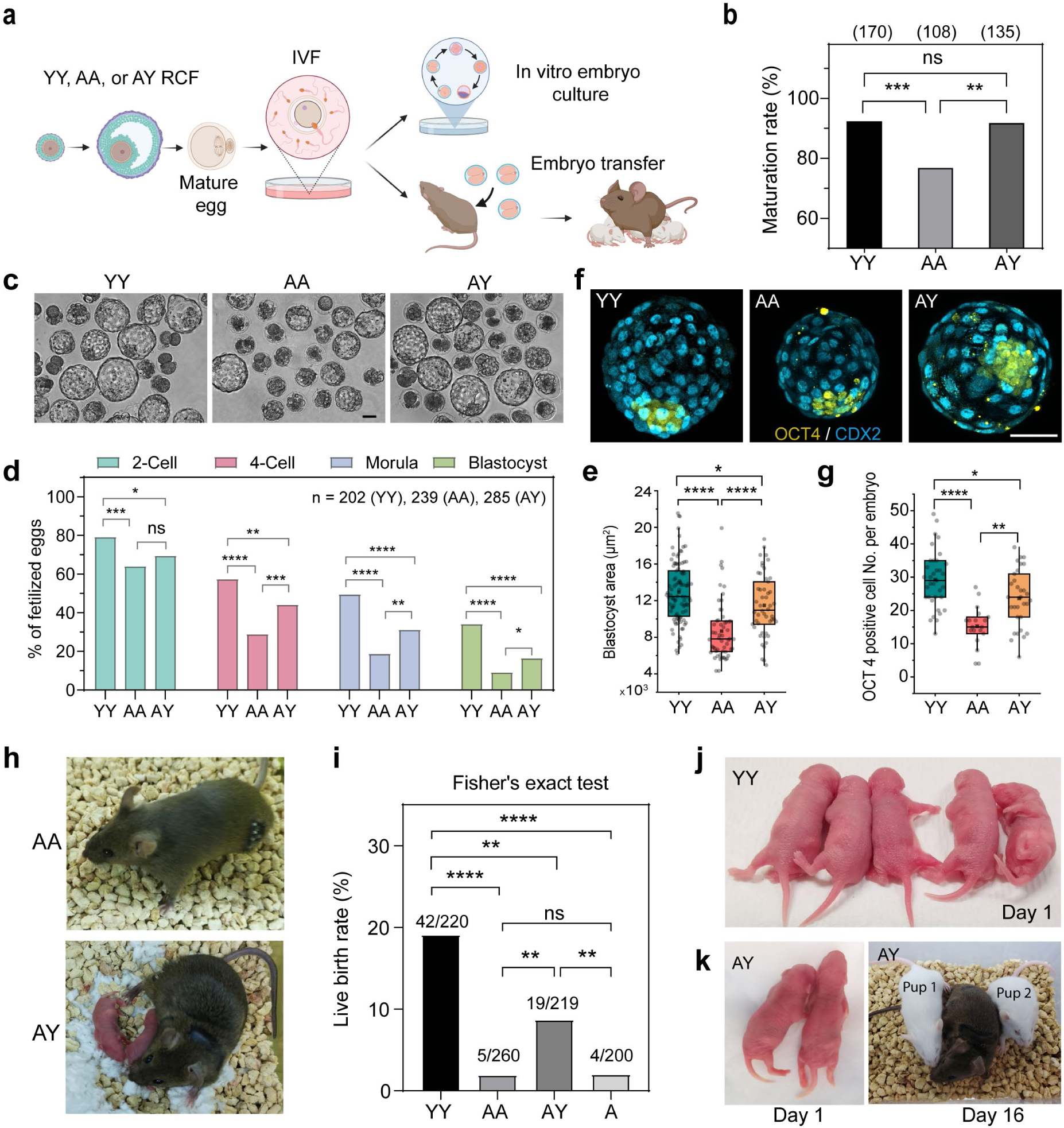
Young r-follicles restored the developmental potential of aged oocytes. **a.** Schematic illustration of the experimental design for assessing embryo development competence. **b.** Comparison of oocyte maturation rates (PB1 extrusion rates) among oocytes from YY, AA, and AY RCFs. The number of oocytes with the PB1 extruded was counted after maturation for 16h. The number of oocytes is specified in brackets. 2-3 month-old (young) and 14-17 month-old (aged) wide type ICR mice were used. Fisher’s exact test. **c.** Representative images of early embryos resulting from YY, AA, and AY RCFs. Scale bars, 50 μm **d.** Analysis of embryo development potential following *in vitro* fertilization (IVF) of oocytes form AA, AY, and YY RCFs. n = 202 (YY), 239 (AA), 285 (AY). 2-3 month-old (young) and 14-15 month-old wide type ICR mice were used. Fisher’s exact test. **e.** Quantification of blastocyst size measured as projected area. n = 99 (YY), 56 (AA), 50 (AY). Box plots indicate mean and median by black square and center bar; upper and lower limits and whiskers represent quartiles and 1.5 × interquartile; one-way ANOVA, Tukey’s multiple comparisons test. 2-3 month-old (young) and 14-15 month-old (aged) wide-type ICR mice were used. **f.** Blastocysts stained with OCT4 (yellow) and CDX2 (cyan) to quantify inner cell mass cells and trophectoderm, displaying representative maximum projection images of blastocysts derived from YY, AA, and AY RCFs. Scale bars, 50 μm. **g.** Quantification of OCT4-positive cells in blastocysts derived from YY, AA, and AY RCFs yield oocytes. n = 35 (YY), 31 (AY), 19 (AA). Box plots indicate mean and median by black square and center bar; upper and lower limits and whiskers represent quartiles and 1.5 × interquartile; one-way ANOVA, Tukey’s multiple comparisons test. 2-3 month-old (young) and 14-15 month-old (aged) wide type ICR mice were used. **h.** Representative images of surrogate mothers and pups from AY and AA RCFs. Note that two pups were delivered after transplanting 20 2-cell stage embryos derived from AY RCFs into a surrogate mother, whereas no pup was delivered after transplanting 20 AA 2-cell stage embryos derived from AA RCFs. **i.** Live birth rates after transferring 2-cell embryos into surrogate mothers (3-6 months old). The numbers in the graph indicate the number of pups born / number of embryos transferred. Follicles from 2-month-old (young) and 14-month-old (aged) wild-type ICR mice were used to generate RCFs. Fisher’s exact test. **j.** Day 1 pups generated from oocytes obtained from YY RCFs. **k.** Day 1 (left) and day 16 (right) pups (white mice) that birthed using oocytes derived from AY RCFs. Note that the surrogate mother had brown fur, while the pups from embryos transferred were of the ICR strain background that exhibits white fur. \*\*\*\**P* < 0.0001; \*\*\**P* < 0.001; \*\**P* < 0.01; \**P* <0.05; ns, P > 0.05.

The embryonic developmental potential of the aged oocytes grown and matured in young follicular environments was assessed after *in vitro* fertilization (Fig. 2c,d). As expected, AA RCFs yielded oocytes that supported a lower rate of blastocyst formation than oocytes from YY RCFs. AY RCFs significantly improved the rate of blastocyst formation of aged oocytes (Fig. 2c,d). The size and total cell number of blastocysts developed from AY RCFs were also significantly increased compared to those from AA RCFs (Fig. 2e, Extended Data Fig. 6e,f).

Improvement was also observed for the development of inner cell mass, as evidenced by an increase in the number of OCT4^+^ cells (Fig. 2f,g), a critical predictor of pregnancy success^28^. The developmental competence of oocytes from different RCFs was further validated by transferring 2-cell embryos to pseudopregnant females. Oocytes from AY RCFs produced a significantly higher (∼3.5x higher) rate of live birth than those from AA RCFs or aged intact follicles (A), although the live birth rate of the former was still lower (∼1x) than those from YY RCFs (Fig. 2h-k). To ensure that the observed improvements were not due to potential defects introduced by the RCF model, we verified that oocytes from aged unperturbed follicles (A) and AA RCFs did not show significant differences in ROS levels, maturation rate, the frequency of spindle abnormalities or chromosome misalignment, or the rate of blastocyst formation or blastocyst size (Extended Data Fig. 7). These results suggest that the young follicular environment can partially restore the quality and developmental competence of aged oocytes.

## Enhanced TZP Formation in AY RCFs

TZPs are a key mediator of GC-oocyte interactions, through which essential nutrients and metabolites are supplied from GCs to the developing oocyte^29^ (Extended Data Fig. 8a). Prior work showed that the quantity of TZPs diminishes during follicle aging^20^, raising the possibility that the improved oocyte quality in AY RCFs may be associated with improved TZP formation. To observe TZP reformation within RCFs, we transplanted wild-type (non-fluorescently labelled) oocytes into r-follicles from the mTmG transgenic mice^30^ (Fig. 3a). Super-resolution microscopy imaging showed the presence of tdTomato-labeled processes, resembling TZP, extending from the granulosa cells to the non-fluorescent oocyte, with characteristic bulbous swellings at the tips (Fig. 3b). These tdTomato-labeled processes formed within 3 hours following follicle reconstitution (Extended Data Fig. 8b).

**Figure 3.**
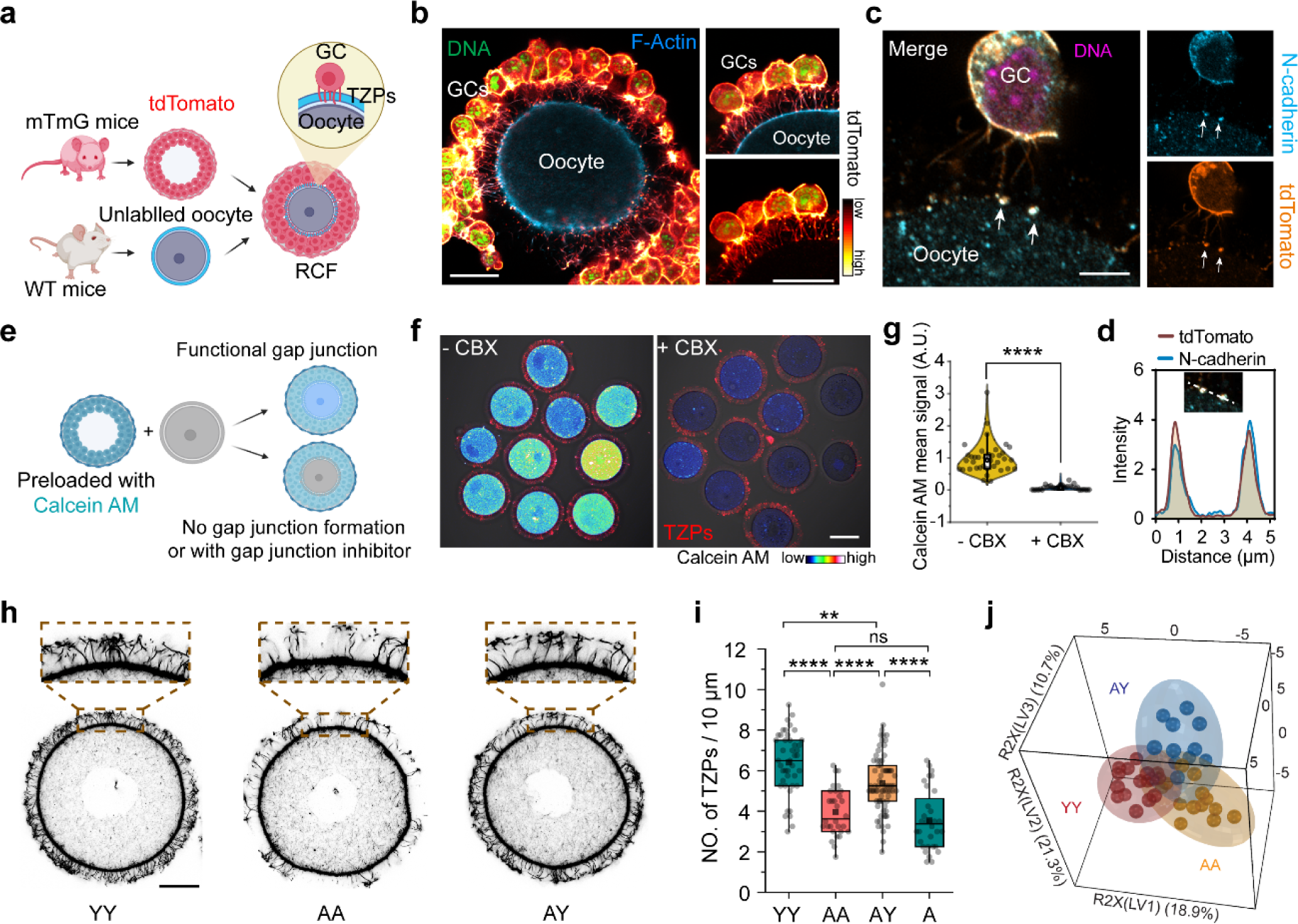
Regeneration of TZP and metabolic remodelling of oocytes in RCFs. **a.** Experimental scheme for monitoring TZP regeneration. To visualize newly generated TZPs, RCFs were created using somatic cells from transgenic mTmG mice expressing membrane-targeted tdTomato and unlabelled oocytes from wild-type mice. **b.** Airyscan super-resolution images of tdTomato-labeled processes extending from mTmG granulosa cells to wild-type oocytes, forming bulbous swellings at the tips after reconstitution. To enhance TZP visibility, somatic cells were partially removed. The oocyte cortex was stained with phalloidin (cyan), while TZPs from mTmG granulosa cells were stained with anti-RFP. DNA (pseudo-colored green) was stained with DAPI. The left panel displays an example of a 3-color (tdTomato, F-actin, DNA) merged image featuring the entire oocyte. The right panels present another example with a cropped view. The top right panel is similar to the left panel but with a cropped view, while the bottom right panel shows only tdTomato and DNA staining. Scale bars, 20 μm. **c.** Airyscan super-resolution image of N-cadherin (cyan) staining. Arrows show N-cadherin foci at the tip of TZP associated with oocyte surface. The left panel displays a 3-color (tdTomato, N-cadherin, DNA) merged image. The top right panel shows only N-cadherin staining, while the bottom right panel shows only tdTomato staining. The RCFs were created using the same method as described in (b). DNA (magenta) was stained with DAPI. Scale bars, 5 μm. **d.** Intensity profiles of N-cadherin and tdTomato along the dashed line. **e.** Experimental design to investigate gap junction communication within RCFs. Follicular somatic cells were pre-incubated with Calcein AM for 1 hour to enable transport of the dye to somatic cells. These cells were then combined with unlabelled wild-type oocytes to create RCFs, which were cultured in the presence or absence of a gap junction inhibitor, carbenoxolone (CBX). Calcein AM signals were observed 3 hours later to determine the extent of dye transfer. **f.** Confocal microscopy images of Calcein AM fluorescence in oocytes from RCFs with and without gap junction inhibitor CBX treatment. Follicular somatic cells were removed, leaving behind only TZPs (tdTomato, red). Color bar shows the intensity of Calcein AM. Scale bars, 50 μm. **g.** Mean fluorescence intensity of Calcein AM in oocytes was quantified. n = 37 (-CBX), 35 (+CBX). Two-tailed unpaired t-test. \*\*\*\**P* < 0.0001. **h,i.** Airyscan super-resolution images of phalloidin-stained TZPs in oocytes from YY, AA, and AY RCFs (**h**). Enlarged views of the areas within the brown dotted rectangles are presented above. Scale bars, 20 μm. (**i**) The density of TZPs was then quantified. n = 43 (YY), 30 (AA), 74 (AY), 32 (A). Experiments utilized 2-month-old (young) and 14-month-old (aged) wild-type ICR mice. Box plots indicate mean and median by black square and center bar; upper and lower limits and whiskers represent quartiles and 1.5 × interquartile; one-way ANOVA, Tukey’s multiple comparisons test. \*\*\*\**P* < 0.0001; \*\**P* < 0.01; ns, P > 0.05. **j.** The PLS-DA 3D score plot (R2X_cum_ = 0.508, R2Y_cum_ = 0.684, Q2_cum_ = 0.445) showing separations of metabolic features among oocytes from AA, YA, and YY RCFs. Ellipses indicated the 95% confidence interval. Each colored spot represents one biological replicate. 2-month-old (young) and 14 month-old (aged) wide type ICR mice were used.

To verify that these are functional TZPs, we examined the presence of adherens junction and gap junction at the interface between the processes and the oocyte, as those are essential features of TZPs (Extended Data Fig. 8a). Previous studies suggested that adhesion depends on the interaction between N-cadherin on the surface of TZPs and E-cadherin on the oocyte surface^16,31^. Indeed, immunofluorescent staining showed the presence of N-cadherin at the tip of the tdTomato-labeled processes that were in contact with the oocyte (Fig. 3c,d). We examined gap junctional communication between the oocyte and granulosa cells by implanting a unlabeled, wild-type oocyte into r-follicles pre-incubated with Calcein AM, a gap-junction-permeable fluorescent dye (Fig. 3e). Calcein AM fluorescence was detected in the oocyte after co-culture, and this dye transfer was prevented when the RCF was incubated with carbenoxolone (CBX), a pharmacological inhibitor of gap junction activity (Fig. 3f,g). These results suggest that GC within RCFs can extend functional TZPs to reach the oocyte surface, re-establishing communication between the oocyte and the surrounding somatic cells. Importantly, AY RCFs formed a greater number of TZPs per unit length of the oocyte periphery than AA RCFs or aged intact follicles (A), although the TZP density was still lower than YY RCFs (Fig. 3h,i).

## Metabolic and Transcriptomic Remodeling of Oocytes in RCFs

To investigate whether young follicular environment can alter the metabolic profile of aged oocytes, we performed a liquid chromatography-mass spectrometry (LC-MS/MS)-based metabolomics analysis. 136 metabolites were identified, with 34 showing differential abundance comparing oocytes from YY, AA, and AY RCFs (Supplementary Table 2). Partial least squares discriminant analysis (PLS-DA) showed that the metabolic profile of oocytes from AY RCFs partially separated from that of oocytes from AA RCFs, suggesting metabolic remodelling of aged oocytes by the young r-follicles (Fig. 3j). Importantly, among the 17 metabolites that differed between YY and AA oocytes, 13 in AY oocytes displayed similar levels to those in YY oocytes (Extended Data Fig. 8c).

We further investigated whether the transcriptome of aged oocytes was also remodeled within AY RCFs by performing single-oocyte RNA sequencing on oocytes from YY, AY, and AA RCFs. Our analysis identified a significant divergence, encompassing 1,774 differentially expressed genes (DEGs) between oocytes from YY and AA RCFs, with 1,116 genes upregulated and 658 downregulated (Extended Data Fig. 9a, Supplementary Table 3). Gene Ontology (GO) analyses revealed that downregulated genes in aged oocytes from AA RCFs contribute to processes, such as protein epigenetic modifications and cellular communication, reflecting a reduced dynamic interplay with their microenvironment, while those genes that were upregulated in aged oocytes from AA RCFs were associated with metabolic processes and mitochondrial functions, likely as a counterbalance to aging-induced metabolic alterations (Extended Data Fig. 9b,c).

Importantly, our data revealed that the transcriptomic landscape of aged oocytes from AY RCF resembled more closely that of young oocytes from YY RCFs than that of aged oocytes from AA RCFs (Fig. 4a-c). Culturing aged oocytes in young RCFs reversed the age-associated expression changes of a large number of genes (Fig. 4b,c, Supplementary Table 4, 5). Gene set enrichment analysis (GSEA) revealed that the AY RCFs potentially restored several affected pathways in aged oocytes (Fig. 4d), such as the estrogen signalling, Ras signalling, and nucleotide excision repair, which are all related to oocyte development and quality control^7,32,33^.

**Figure 4.**
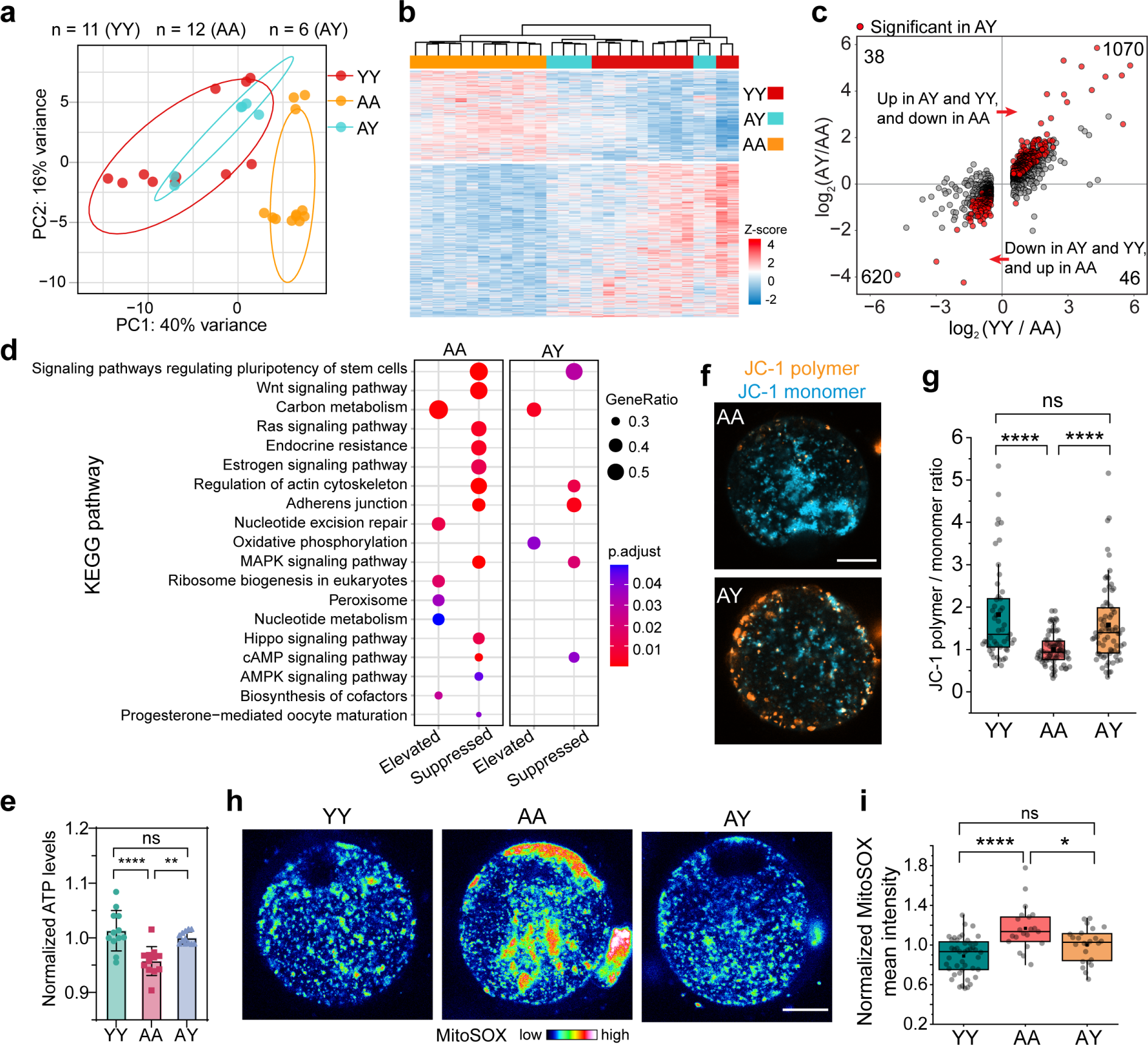
Transcriptomic remodelling and mitochondrial fitness in aged oocytes cultured in young r-follicles. **a.** Principal component analysis (PCA) of the normalized gene expression data in single oocytes from AA, YA, and YY RCFs. Each dot represents a single oocyte from each group. N = 11 (YY), 12 (AA), 6 (AY). Ellipses fit a multivariate t-distribution at confidence level of 0.8. 2-month-old (young) and 14-month-old (aged) wide type ICR mice were used for single oocyte RNA-seq. **b,c.** Heatmap (b) and scatter plot (c) showing mRNA levels of 1,774 genes differentially expressed between young and aged oocytes from YY and AA RCFs, respectively. The corresponding expression of these genes in oocytes from AY RCFs is also shown. The heatmap, generated via unsupervised hierarchical clustering, shows that oocytes in AY RCFs exhibit a shift in gene expression patterns towards the direction of the young oocytes from YY RCFs. The scatter plot reveals that among 1116 upregulated and 658 downregulated genes comparing oocytes from YY with those from AA RCFs, 1070 genes were upregulated and 620 genes were downregulated in oocytes from AY RCFs compared to AA RCFs, respectively. Red dots in (c) represent the genes (245 out of 1070, and 160 out of 620) with significantly changed expression (*P* < 0.05) in AY oocytes compared to AA oocytes. **d.** Dot plot of filtered enriched KEGG pathways from GSEA with each group comparing against oocytes from YY RCFs. Elevated indicates positive normalized enrichment score while supressed indicates negative score. **e.** Quantification of ATP levels in oocytes from YY, AA, and AY RCFs. 2-month-old (young) and 14-month-old (aged) wide type ICR mice were used. Data are shown as mean ± SD. **f,g.** Mitochondrial membrane potential was assessed using JC-1 staining (**f**). Scale bar, 20 µm. The ratio of JC-1 polymer to JC-1 monomer fluorescence intensity (**g**) was calculated for oocytes from YY, AA, and AY RCFs to determine differences in mitochondrial membrane potential. n = 49 (YY), 74 (AY), 72 (AA). 2-month-old (young) and 14-15 month-old (aged) wide type ICR mice were used. Box plots indicate mean and median by black square and center bar; upper and lower limits and whiskers represent quartiles and 1.5 × interquartile. **h.** MitoSOX staining, reporting mitochondrial ROS levels, in MII oocytes from YY, AA, and AY RCFs. Scale bar, 20 µm. **i.** The fluorescence intensity of MitoSOX was measured and compared between oocytes from YY, AA, and AY RCFs. n = 44 (YY), 21 (AA), 24 (AY). 2-month-old (young) and 14-15 month-old (aged) wide type ICR mice were used. Box plots display data distribution, with the mean and median represented by a black square and center bar, respectively; the upper and lower limits, as well as whiskers, correspond to quartiles and 1.5 × the interquartile range. The data in (**e**), (**g**), and (**i**) were analyzed by one-way ANOVA, Tukey’s multiple comparisons test. \*\*\*\**P* < 0.0001; \*\**P* < 0.01; \**P* <0.05; ns, *P* > 0.05.

## Young Follicular Environment Improves Mitochondrial Function of Aged Oocytes

As metabolomic and transcriptomic analyses revealed potential changes in the metabolism and physiology of oocytes from AY RCFs, we further looked into mitochondrial function and fitness, which are critical to oocyte growth, maturation and developmental potential^34^. Indeed, while lower ATP production was observed in aged oocytes from AA RCFs, ATP production was largely restored in oocytes from AY RCFs (Fig. 4e). Confocal microscopic analyses of oocytes stained with MitoTracker revealed that mitochondria in oocytes from AA RCFs exhibited heterogeneous clustered distribution, which was previously shown to be linked to the decline in oocyte quality with advancing age^35^. Mitochondria in oocytes from AY RCFs exhibited more uniform cytoplasmic distribution, reassembly that in YY oocytes, than in oocytes from AA RCFs (Extended Data Fig. 9d,e). Consistent with improved mitochondrial fitness, culturing within young r-follicles largely restored mitochondrial membrane potential in aged oocytes (Fig. 4f,g, Extended Data Fig. 9f). Oocytes from AY RCFs exhibited lower mitochondrial and cellular ROS than aged oocytes from AA RCFs (Fig. 4h,i, Extended Data Fig. 9g,h).

As previous research suggested that mitochondrial transfer from healthy sources could elevate oocyte quality in both aged mice and humans^36–38^, we assessed whether there was mitochondrial transfer from surrounding granulosa cells to oocytes in RCFs. Wild-type (non-fluorescently labelled) oocytes were transplanted into r-follicles from transgenic mice in which mitochondria-targeting sequence (MTS)-mCherry-GFP_1-10_ was expressed from the Rosa26 locus (this mouse line was generated for another study and GFP_1-10_ was irrelevant here). The mitochondrial mCherry signal enabled tracking of possible mitochondrial transfer from GCs to oocyte; however, we did not observe any mCherry-labelled mitochondria within the oocytes transplanted into MTS-mCherry-GFP_1-10_ r-follicles (Extended Data Fig. 10). It was thus unlikely that young r-follicles improved oocyte quality through mitochondrial transport from granulosa cells to oocytes.

## Improved Meiotic Chromosomal Segregation in AY RCFs

Aneuploidy is the main cause of infertility in aged females^39^. To assess chromosome numbers, we performed super-resolution imaging of live MII oocytes injected with mRNA coding for mCherry-tagged histone H2B (H2B-mCherry) and two tandem mEGFP-tagged CENP-C (2mEGFP-CENP-C) mRNA to label chromosomes and kinetochores separately^40^. Subsequent 3D reconstructions of chromosomes and kinetochores enabled accurate visualization and quantification of chromosome and kinetochore numbers in oocytes (Fig. 5a). We quantified “chromosomal abnormalities,” which include both altered numbers of sister kinetochores (aneuploidy) and premature separation of sister chromatids (PSSC). A significantly higher frequency of chromosomal abnormalities was observed in aged oocytes from AA RCFs compared to young oocytes from YY RCFs (Fig. 5a,b), which is consistent with *in vivo* studies^41,42^. Importantly, oocytes from AY RCFs exhibited a reduced rate of chromosomal abnormalities compared to oocytes from AA RCFs (Fig. 5b).

**Fig. 5.**
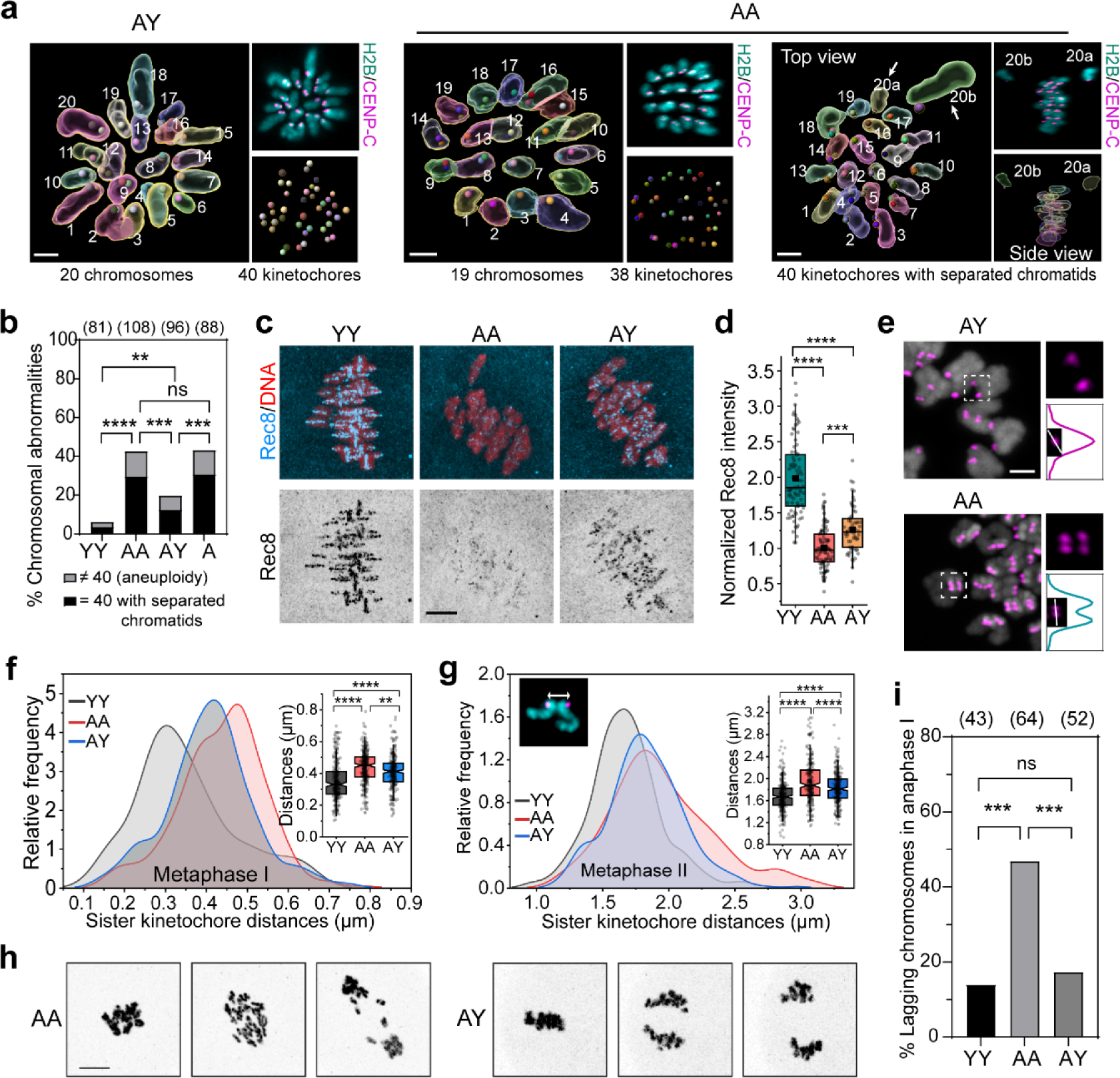
Improved meiotic fidelity in aged oocytes cultured in young follicular environment. **a.** 3D reconstruction of MII chromosomes and kinetochores for assessment of chromosome abnormalities. Oocytes expressed 2mEGFP-CENP-C and H2B-mCherry to label kinetochores and chromosomes, respectively. AY: The left panel displays the 3D reconstructed chromosomes and kinetochores, with chromosome numbers labelled. The top-right panel shows Z-projection images of 2mEGFP-CENP-C and H2B-mCherry, while the bottom-right panel displays 3D-reconstructed kinetochores. In the representative image, a total of 20 chromosomes with 40 kinetochores (euploidy) were found. AA (Left): the panels display the same as AY, but in the representative image, a total of 19 chromosomes and 38 kinetochores (aneuploidy) were found. AA (right): the left panel displays the top view of the metaphase plate with 3D reconstructed chromosomes and kinetochores, with chromosome numbers labelled. Arrows indicate separated chromatids. The side view of the metaphase plate with 3D reconstructed chromosomes and kinetochores is shown in the bottom-right panel, while the corresponding Z-projection image of 2mEGFP-CENP-C and H2B-mCherry is shown on top-right panel. Note that in the representative image, a total of 40 kinetochores were found but with the separated chromatids. Images were generated using Imaris software. Scale bar, 2 µm. **b.** Quantification of the percentage of chromosomal abnormalities in oocytes from YY, AA, and AY RCFs, and from aged intact follicles (A). n = 81 (YY), 108 (AA), 96 (AY), 88 (A). “Chromosomal abnormalities” include: 1) Number of kinetochores ≠ 40 (aneuploidy), and 2) Number of kinetochores = 40 with separated chromatids. 2-month-old (young) and 14-month-old (aged) wide-type ICR mice were used. The data was analyzed by using Fisher’s exact test. **c,d.** Young follicular somatic cells partially restore the aging-associated reduction of chromosomal cohesin. (**c**) Oocytes were stained for Rec8 (cyan) and DNA (Hoechst, red) at MI stage. Z-projection images are shown. Scale bar, 5 μm. Chromosomal Rec8 signals were quantified as shown in (**d**). Box plots indicate mean and median by black square and center bar; upper and lower limits and whiskers represent quartiles and 1.5 × interquartile; one-way ANOVA, Tukey’s multiple comparisons test. 2-month-old (young) and 14-month-old (aged) wide-type ICR mice were used. **e.** Maximum intensity z-projection immunofluorescence images of kinetochores (2mEGFP-CENP-C) and chromosomes (H2B-mCherry) in MI oocytes from AA and AY RCFs. Top-right images are magnifications of outlined regions, while bottom-right images display intensity profiles of a sister kinetochore pair along the white line in the insets. Scale bar, 2 µm. **f.** Sister kinetochore pair distances for MI oocytes from YY, AA, and AY RCFs were measured. See the Methods section for details on distance measurement for MI oocytes. Kernel-smoothed distribution curves are displayed. The inset notched boxplot represents the distances of sister kinetochore pairs for MI oocytes from YY, AA, and AY RCFs. n = 278 sister kinetochore pairs for YY RCFs, 305 for AA RCFs, and 265 for AY RCFs. **g.** Sister kinetochore pair distances for MII oocytes were measured. The relative frequency of individual sister kinetochore pair distances for YY, AA, and AY groups is presented, with kernel-smoothed distribution curves. The left inset displays a paired set of sister chromatids and kinetochore. The inset notched boxplot represents the distances of sister kinetochore pairs for MII oocytes from YY, AA, and AY RCFs. n = 299 sister kinetochore pairs for YY RCFs, 272 for AA RCFs, and 292 for AY RCFs. See Methods and Supplementary Video 2 for details on the measurement protocol. **h.** Representative time-lapse frames from high-resolution live-cell imaging of meiosis I division in oocytes, with chromosomes (grey) labelled with H2B-mCherry. Scale bar, 10 µm. **i.** Quantification of the percentage of lagging chromosomes observed in the experiments in (**h**). n = 43 (YY), 64 (AA), 52 (AY). The data was analyzed by using Fisher’s exact test. 2-month-old (young) and 14-month-old (aged) wide-type ICR mice were used. Notched boxplots in (**f**) and (**g**) illustrate data distribution, with boxes marking medians and quartiles, notches indicating a 95% confidence interval for the median, and whiskers extending to 1.5× the interquartile range. The data in (**d**), (**f**), and **(g**) was analyzed by using one-way ANOVA, Tukey’s multiple comparisons test. ****P < 0.0001; ***P < 0.001; **P < 0.01; ns, P > 0.05.

Age-related aneuploidy in oocytes is in part attributed to the premature loss of cohesin prior to meiotic divisions^43–45^. We therefore performed immunofluorescence staining of Rec8, a meiotic cohesin component, and measured its signal intensity on chromosomes (Fig. 5c). The results show that the level of chromosomal Rec8 in aged oocytes from AA RCFs were significantly reduced compared to that in oocytes from YY RCFs, consistent with age-associated cohesin loss in previous studies^43^. Notably, the level of chromosomal Rec8 in oocytes from AY RCFs was significantly higher than that in AA RCFs, albeit not as high as that in YY RCFs (Fig. 5c,d). We then measured the distance between paired (excluding PSSC) sister kinetochores at MI and MII, which can be regarded as a measure of centromere cohesion between sister chromatids, with larger distances indicating less cohesion^43–46^. The results showed that oocytes from AY exhibited shorter distances between sister kinetochores than oocytes from AA RCFs (Fig. 5e-g, Extended Data Fig. 11a, Supplementary Video 2), suggesting that the young follicular microenvironment helped maintain centromere cohesion in aged oocytes.

Spindle abnormalities and lagging chromosomes at anaphase are another cause of aneuploidy ^41^. To monitor chromosome segregation, oocytes were microinjected with H2B-mCherry to label chromosomes and then subjected to 3D time-lapse imaging (Fig. 5h). As expected, the incidence of lagging chromosomes during the anaphase of meiosis I was significantly lower in oocytes from AY RCFs than AA RCFs (Fig. 5i). Consistent with a higher occurrence of lagging chromosomes, aged oocytes from AA RCFs contained more incorrect kinetochore-microtubule (KT-MT) attachments (unattached, merotelic, and lateral attachment), compared to young oocytes from YY RCFs (Extended Data Fig. 11b-d). The rate of incorrect KT-MT attachments was significantly decreased in the aged oocytes cultured with young r-follicles (Extended Data Fig. 11b-d). Taken together, these data show that the young follicular microenvironment improved meiotic chromosome transmission in oocytes through better maintenance of chromosome cohesion and reduction in chromosome mis-segregation.

## Discussion

By utilizing the mouse ovarian follicle as a model, the study presented above underscores the significant impact of the aging follicular microenvironment on the developmental potential of oocytes. By constructing chimeric follicles, we showed that an aged follicular environment could harm the quality and developmental potential of young oocytes. More remarkably, a young follicular environment could restore the maturation efficiency of oocytes from older animals and enhance subsequent embryonic development and live birth rates.

The observed improvements in the quality of aged oocytes was likely to be mediated by improved communication between oocytes and surrounding somatic cells through TZPs. Gap junctions at the TZP-oocyte interface permit the transfer of essential metabolites, nucleotides, amino acids and ions from granulosa cells to the oocyte to support oocyte growth and development. The more youthful GCs and their superior proficiency in TZP formation could lead to enhanced metabolic flux toward oocyte in the AY RCFs (Extended Data Fig. 12). Previous work showed that gap junction-mediated GC-oocyte communication plays an important role in influencing oocyte transcriptional activities, chromatin organization, and post-translational modifications of oocyte proteins^47–49^. As such, the enhanced GC-oocyte communication and metabolic flux could lead to metabolic and transcriptomic remodelling in the aged oocyte and mitigate mitochondrial dysfunction. The resulting increased ATP production could better support the energy-expensive processes, such as meiotic chromosome segregation and polar body extrusion, and the reduced ROS could underlie improved preservation of cohesion and fidelity of chromosome segregation.

Our data shows the AY RCF culture partially restored Rec8 cohesin level, which help explain the improved centromere cohesion and chromosome segregation in the aged oocytes. Evidence from mouse oocytes suggests that the loading of cohesin complexes onto chromosomes is limited to fetal development^50^. Thus, the observed partial recovery of cohesin in AY RCFs is likely to reflect a reduction in cohesion loss during the active follicle growth from the secondary to the antral stage, as well as during oocyte maturation. One possible explanation is the reduced oxidative stress in the aged oocyte in AY RCFs, as evidenced by the reduced ROS. This could be due to the young granulosa cells supplying antioxidants and other protective factors to the oocyte^51,52^ or due to improved mitochondrial function in the oocyte. Oxidative stress can also lead to DNA damage, protein modifications, and perturbed proteostasis, all of which could adversely impact cohesin stability. Moreover, our RNA-seq data indicates that Sgo2b, a gene associated with cohesin protection^44^, was upregulated in oocytes from AY RCFs, potentially mitigating cohesin loss.

It is worth noting that the observed rejuvenation of aged oocytes by young follicular cells was incomplete, as blastocyst formation and live birth rates associated with oocytes from AY RCFs were still lower than those from YY RCFs. Aged oocytes may endure changes, such as nuclear or mitochondrial genome damage, extensive cohesin loss, epigenetic modifications, or proteostasis loss^7,53–56^, that cannot be fully repaired or fully reversed. Another possible explanation for the partial rejuvenation is the presence of the reverse signalling from aged oocytes to the surrounding young somatic cells in AY RCFs. Granulosa cells are known to rely on the oocyte for growth factors such as GDF9 and BMP15^57,58^. Although GDF9 was supplemented in the culture media for RCFs, other factors might be insufficient from the aged oocyte or supplied insufficiently from the media. Aged oocytes might also release factors that negatively affect the function and quality of the young somatic cells. Further research is needed to identify the potential limiting or toxic factor(s) involved in order to maximize the rejuvenation potential of our approach. Despite the less than full rejuvenation, the combination of the fully restored rate of oocyte maturation and the partially restored embryonic development to-term, a somatic cell-based rejuvenation therapy prior to IVF, as suggested by our findings, could considerably enhance the change of viable offspring.

In summary, our finding has significant clinical implications, as it suggests that the age-related decline in oocyte quality can potentially be reversed by co-culturing with young and healthy somatic cells and possible also the associated extracellular environment present in young follicles, providing the basis for a potentially safe cell-based therapeutic strategy to treat age-associated female infertility. Establishment of this therapy will require further identifying the minimum follicular components required for the rejuvenation and a method of reconstituting these components with oocytes *in vitro*. In addition, the RCF culture developed in this study provides a model system for dissecting age-related changes that may be differentially attributed to oocytes and the surrounding somatic cell environment and mechanisms underlying these changes.

## Acknowledgments

We thank Shuo Xiao (Rutgers University) for the helpful discussion on mouse follicle in vitro culture method. We thank Michael Lampson for providing Rec8 antibody. We thank Tomoya S. Kitajima (RIKEN Center for Developmental Biology) for providing pGEMHE-2mEGFP-CENP-C plasmid. We thank Andrew Ewald (Johns Hopkins University School of Medicine) for providing mTmG mice strain. This work was supported by grant from NUS Bia-Echo Asia Centre for Reproductive Longevity and Equality (ACRLE).

## Author contributions

H.W., H.Z., and R.L. conceived the study. H.W. and R.L. designed the experiments and methods for data analysis. H.W. performed experiments and analyzed the data with assistance from X.J.S. and X.S., with the following exceptions: L.H.W., P.L.L., and L.S.P. performed the MS experiments and data analysis; C.S. measured the distance between sister kinetochores; X.Z. generated MTS-mCherry-GFP_1-10_ mice strain; J.Z. supervised the RNA-seq experiments and analyzed the data with Y.L.; and C.L.D. and L.S.P. supervised the MS analysis. H.W. and R.L. wrote the manuscript and prepared the figures with input from all authors. R.L. supervised the study.

## Additional Information

Correspondence and requests for materials should be addressed to Rong Li or HaiYang Wang.

## Methods

### Animals

The animal experiments conducted in this study were approved by the Institutional Animal Care and Use Committee at the National University of Singapore and were carried out in compliance with the relevant guidelines and regulations. The mice were kept under standard conditions, with constant temperature (21-25°C) and humidity (30-70%), and a 12-hour light:12-hour dark cycle with light onset at 07:00. The mice utilized in our experiments were kept as virgin mice throughout their lifespan. ICR mice were procured from Singapore InVivos, while mTmG mice were obtained as a gift from the Andrew Ewald lab at Johns Hopkins University.We used mTmG transgenic mouse strain specifically in Fig 1i, Fig 3a-g, and Extended Data Fig. 5. All comparisons between groups YY, AY, AA, and YA were conducted with wild-type ICR mice.

Parental mice C57BL/6J for making the MTS-mCherry-GFP_1-10_ transgenic mice were obtained from the Jackson Laboratories (strain #000664). The MTS-mCherry-GFP_1-10_ transgenic mice were generated by Transgenic Core Laboratory in Johns Hopkins University, using homology-directed repair based, CRISPR-Cas9-induced precise gene editing^59^. CAG promoter driven MTS (Su9)-mCherry-GFP_1-10_ sequence was inserted into the safe harbor *Rosa26* locus using plasmid containing the construct flanked by *ROSA26* homologous sequence (1083 bp upstream/4341bp downstream overlap). They were microinjected together with CRISPR Cas9 protein and crRNA (CGCCCATCTTCTAGAAAGAC) into one-cell embryos of C57BL/6 *WT* mice. PCR genotyping was performed with primer pairs forward TTCCCTCGTGATCTGCAACTC and reverse CTTTAAGCCTGCCCAGAAGACT for *WT Rosa*26; forward GTGGGAGCGGGTAATGAACTTT and reverse TCCTGCAATGATGAATCTTGAGTGA for MTS-mCherry-GFP_1-10_ knock-in. The correct insertion generated a band size of 69 bp. Homozygous MTS-mCherry-GFP_1-10_ mice were PCR genotyped and used in this study.

### Preparation of Alginate-rBM IPN

To prepare sterile alginate solution, alginate powder (#71238, Sigma-Aldrich) was dissolved in MilliQ H_2_O with continuous stirring overnight to a final concentration of 1% w/v. Activated charcoal was added at a ratio of 0.5 g per gram of alginate, and the mixture was stirred for 30 minutes. The alginate solution was then sterile-filtered through Millipore Express 0.22 μm filters and transferred to a 50 mL sterile tube. The alginate solution was frozen at −20°C for 1-2 days and then lyophilized for approximately 5 days until dryness. The dried alginate was stored at −80 °C until needed. Before use, the alginate was reconstituted with 1× PBS without Ca^2+^ to the concentration of 1% w/v as a stock solution. The mixture was vortexed briefly and then left on an orbital shaker until it was completely dissolved.

Matrigel (#354230; Corning) was purchased for use as the rBM matrix. Alginate-rBM IPN was prepared by mixing the alginate stock solution with growth media (to achieve a final concentration of 0.25% w/v), Matrigel (at a final concentration of 0.1 mg/ml) on ice. A 4.5 µL Alginate-rBM IPN bead was then crosslinked in a solution composed of 50 mM CaCl_2_ and 140 mM NaCl for 3-4 min.

### Follicle isolation, encapsulation, and culture

Ovaries were dissected from 2-3 month-old (young) or 14-17 month-old (aged) virgin female mice. Individual follicles were obtained by breaking down the ovaries into small pieces with a 26 G needle and incubating them in Dissection Media containing L-15 (Thermo Scientific, #11415064) with 1% fetal bovine serum (FBS) and 100 U/ml penicillin/streptomycin supplemented with 2 mg/ml collagenase (SCR103, Sigma) and 10 U/ml DNAse (D4263, Sigma) for 30-40 minutes. Secondary follicles were then isolated and placed in Maintenance Media containing αMEM (Glutamax, Thermo Scientific #32561102) with 5% FBS and 100 U/ml penicillin-streptomycin for 2 hours before encapsulation. Each follicle was individually encapsulated in a 4.5 ul drop of Alginate-rBM IPN bead. Following a single wash with Maintenance Media, the IPN beads were placed in either 96-well or 24-well plates. Follicles were then cultured in Growth Media, composed of a 1:1 mixture of αMEM Glutamax and F-12 Glutamax, enriched with 5% FBS, 100 mIU/ml follicle-stimulating hormone (FSH, Sigma), 5 μg/ml insulin, 5 μg/ml transferrin, and 5 μg/ml selenium. The follicle culture was maintained at 37°C, with half of the growth media refreshed every other day.

To harvest sufficient number of follicles, multiple same-aged mice were used at the same time, and their follicles were pooled together. These pooled follicles were then equally distributed between AY and AA RCFs for each experiment to avoid potential biases or variations due to age disparities or individual mouse differences.

### Generation of Reconstituted Chimeric Follicle (RCF)

The isolated secondary follicles were first cultured 2 days in Alginate-rBM IPN hydrogel to facilitate the RCF process. Post-incubation, oocytes were gently denuded from cultured secondary follicles using repeated mouth pipetting with a fine oocyte-sized glass pipette. Once denuded, the oocytes were immediately designated for RCF production as donor oocytes.

To generate RCFs, a 27G needle tip was bent by gently dragging it across a petri dish surface. First, use this bent needle tip to anchor and stabilize the r-follicles designated for transplantation, preventing unwanted movement during the RCFs formation process. Subsequently, gently pierce the follicle with a fine oocyte-sized glass mouth pipette and carefully aspirate the oocyte, leaving an empty oocyte pocket within the follicle. Promptly pick up a denuded oocyte using the same fine oocyte-sized glass mouth pipette and gently release it into the follicle pocket. Continue with the same process for the next follicle. After generating 3-10 RCFs, they were immediately encapsulated in Alg-rBM IPN beads and cultured in Growth Media supplemented with 100 ng/ml GDF9. The process is repeated to generate another set of 3-10 RCFs. On the following day, half of the media was replaced with fresh Growth Media without GDF9. The RCFs were then cultured for an additional 4-5 days, reaching a total culture period of 5-6 days, with half of the media being refreshed every other day.

### Oocyte maturation, fertilization, and embryo culture

Follicles were extracted from the Alg-rBM IPN beads using 10 IU/mL of alginate lyase (A1603, Sigma). The oocytes were then matured in Maturation Medium (αMEM with 10% FBS, 1.5 IU/ml human chorionic gonadotropin [hCG], 10 ng/ml epidermal growth factor [EGF], and 100 mIU/ml FSH) for 16 hours at 37°C in 5% CO2 in air.

Oocytes matured *in vitro* were used for *in vitro* fertilization (IVF). To prepare for IVF, caudae epididymides from two 3-4 month old ICR male mice were lanced in 2 drops (100μL / drop) of FERTIUP® Preincubation medium (KYD-002-05-EX, Cosmo Bio) under mineral oil to release sperm, followed by capacitation for 1 hour at 37°C and 5% CO_2_. MII oocytes were then placed in 100 μL of mHTF medium (KYD-008-02-EX-X5, Cosmo Bio) for 30 minutes at 37°C and 5% CO_2_ before being fertilized with 3 μL of sperm suspension. After 6 hours of fertilization, wash the zygotes 3 times with mHTF and culture them in 50 μl mHTF overnight. Next, transfer the resulting 2-cell stage embryos to Continuous Single Culture Media (CSCM-C, Fujifilm #90165) and culture them for 4 days at 37°C, or transfer them to pseudo-pregnant female mice.

### Embryo transfer

Female B6C3HF2 mice (3-6 months old, brown fur) were employed as surrogate mothers in this study. Recipient female (0.5 dpc Pseudo pregnant mouse) was anesthetized with 2.5% Avertin. Animal was checked for loss of pedal reflex and sprayed down with ethanol. A small incision is then made along the dorsal midline of the skin and muscle layer of the left side of the animal. A drop of Epinephrine was placed on the muscle layer prior to incision to prevent excessive bleeding. The ovarian fat pad was seized with forceps and pulled through the incision, carrying with it the ovary, the oviduct, and the upper part of the left uterine horn. Under a stereomicroscope with optic fiber light, a tear is made at the ovarian bursa to expose the infundibulum of oviduct and prepare for oviduct transfer. Tip of Transfer pipette (loaded with HEPES-buffered media and embryos flanked by ‘Air bubbles’) is inserted into the infundibulum and embryos are blown into the oviduct with a mouth pipette. Air bubbles within the ampulla indicate successful transfer. Uterus, oviduct, and ovary are replaced back inside the body cavity. The muscle and skin layer are sutured. Earlier steps are repeated to transfer additional embryos to the right oviduct. Each recipient female was implanted with 18-20 two-cell stage embryos (ICR mice strain). On day 16 post-birth, the fur color of the pups was examined. The surrogate mother mice possessed a brown fur phenotype, contrasting with the white fur exhibited by the ICR strain embryos used. Hence, fur color served as a distinguishing marker between pups originating from donor embryos and potential progeny of the surrogate mother.

### Transcription and oocyte microinjection

Capped mRNA was synthesized from linearized plasmid templates using the T7 or T3 mMESSAGE mMACHINE Transcription Kit (Ambion), polyadenylated (Poly(A) Tailing Kit, Ambion), and purified with the RNeasy MinElute Cleanup Kit (QIAGEN). Approximately 5 to 10 pl of capped mRNAs were microinjected into the cytoplasm of oocytes using a micromanipulator (IM-300 microinjector, Narishige). After injection, the oocytes were cultured at 37°C with 5% CO2 in M2 medium supplemented with 0.2 mM IBMX for at least 3 hours to allow protein expression. Oocytes were microinjected with the following RNAs: H_2_B-mCherry^60,61^, 2mEGFP-CENP-C (a gift from Tomoya Kitajima lab).

### Confocal microscopy

For live follicles, images were acquired with a Zeiss LSM710 confocal microscope with 10x or 20x objectives. For live oocytes, images were acquired with Zeiss LSM980 microscope using a 40× C-Apochromat 1.2–numerical aperture water-immersion objective. Follicles and oocytes were maintained at 37°C with 5% CO2 during imaging. For fixed follicles or oocytes, images were acquired using the Zeiss LSM980 microscopes and processed after acquisition using ZEN (Zeiss).

### Probes for live cell imaging

Live oocytes or follicles were stained with different dyes for confocal microscopy analysis. For mitochondrial staining, oocytes were stained with 200 nM MitoTracker Red CMXRos (M7512, Invitrogen) for 40 min. To measure mitochondrial membrane potential, oocytes or follicles were stained with 2 μM JC-1^62–64^ (M34152, Invitrogen) or 25 nM of TMRM with 100 nM of MitoTracker Green (M7514, Invitrogen). Intracellular ROS was measured by incubating oocytes or follicles with 5 µM CM-H2DCFDA (C6827, Invitrogen). Mitochondrial ROS was measured by incubating oocytes with 500 nM MitoSOX Red (M36008, Invitrogen). To detect apoptosis, follicles were incubated with 500 nM CellEvent™ Caspase-3/7 Green Detection Reagent (C10427, Invitrogen). Oocytes were stained with 150 nM Sir-Tubulin (CY-SC002, Cytoskeleton) and Hoechst 33342 to visualize spindle and chromosomes. All the probes were stained at 37°C in a dark environment with 5% CO2.

### Immunofluorescence

Oocytes, follicles, or embryos were fixed in 4% paraformaldehyde in phosphate-buffered saline (PBS) for 40 minutes and then incubated in a membrane permeabilization solution (0.5% Triton X-100) for 40 minutes. After overnight blocking with 10% bovine serum albumin in PBS, the samples were incubated with the primary antibody overnight at 4°C, followed by incubation with a secondary antibody at room temperature for 1-3 hours. Primary antibodies used were rabbit anti–Ki-67 (#9129S, CST; 1:100), mouse anti-Oct3/4 (sc-5279, Santa Cruz Biotechnology; 1:50), rabbit anti–CDX2 (NB100-2136, Novus Biologicals; 1:100), rabbit anti– RFP (# 600-401-379, Rockland; 1:100), chicken anti–RFP (# 600-901-379, Rockland; 1:100), rabbit anti–Phospho-Histone H2A.X (Ser139) (#9718, CST; 1:100). human anti-Centromere Protein Antibody (ACA, #15-234, Antibodies Incorporated; 1:50), rabbit anti-N Cadherin antibody (ab18203, Abcam; 1:100), rabbit anti-Rec8 antibody^43^ (a gift from Lampson Lab). For secondary antibodies, Alexa Fluor 488–labeled anti-mouse (Invitrogen), Alexa Fluor 488– labeled anti-rabbit (Invitrogen), Alexa Fluor 568–labeled anti-mouse (Invitrogen), Alexa Fluor 568–labeled anti-rabbit (Invitrogen), Alexa Fluor 568–labeled anti-chicken (Invitrogen), Alexa Fluor Plus 647–labeled anti-mouse (Invitrogen), Alexa Fluor Plus 647–labeled anti-rabbit (Invitrogen), Alexa Fluor Plus 647–labeled anti-human (Invitrogen) antibodies were used. F-actin was stained with Alexa Fluor 488–labeled phalloidin (#8878, Cell Signaling Technology) or Alexa Fluor 555–labeled phalloidin (# 8953, Cell Signaling Technology). DNA was stained with DAPI or Hoechst 33342.

### Analysis of gap junctional commutations

R-follicles were incubated for 1 hour in Maintenance Media containing freshly prepared 2 µM Calcein-AM (C3100MP, Invitrogen). Subsequently, oocytes without Calcein-AM labeling were transplanted into Calcein-AM labeled r-follicles to generate RCFs. The aRCFs were then encapsulated in Alg-rBM IPN and cultured either with or without the gap junction blocker Carbenoxolone (ab143590, Abcam) for 3 hours. Following this, the hydrogels were removed, and the denuded oocytes were immediately imaged using a Zeiss LSM980 microscope.

### Single-cell RNA sequencing and data analysis

To obtain pure oocyte samples, devoid of any contamination from surrounding somatic cells, we used a standard oocyte isolation protocol by using precision mouth pipetting techniques. This ensures the thorough removal of any associated granulosa cells. After the isolation, we rigorously inspected the oocytes under the microscope to verify the complete absence of any adhering cumulus cells. To further ensure oocyte purity, an additional expert team member independently verified each oocyte.

Single oocytes were collected with NEBNext Single Cell Lysis Module (NEB #E5530S) in 8μl volume and flash-frozen and stored at −80°C. NEBNext Single Cell/Low Input RNA Library Prep Kit (NEB #E6420) was used for cDNA synthesis, amplification and library generation. Manufacturer’s protocol for cells was followed with these specifications: 16 PCR cycles for cDNA amplification, 8 PCR cycles for enrichment of adaptor-ligated DNA with unique dual index primer pairs from NEB. The libraries were sequenced on the NovaSeq 6000 system with NovogeneAIT Genomics. Reads were aligned to reference mm39 with STAR (2.7.8a) and quantified with HTSeq (0.11.0) using PartekFlow software (v10.0.22.1005) with default parameters.

Differential gene expression analysis was conducted with DESeq2 (Bioconductor 3.16)^65^. Regularized logarithm transformed count data were used for PCA and distance matrix calculation to see the similarities and dissimilarities between samples. To generate the heatmap displaying individual gene expression differences, the subset of normalized count data was further Z-score normalized by row. The DEGs were defined by applying thresholds of absolute log_2_ FC > 0.5 and p.adjust < 0.05. Heatmap with unsupervised hierarchical clustering was generated with the pheatmap package (CRAN 1.0.12) and Volcano plots were generated with EnhancedVolcano (Bioconductor 3.16). Gene Set Enrichment Analysis was carried out using clusterProfiler (Bioconductor 3.16)^66,67^.

### LC/MS-MS assays

Formic acid, Ammonium Formate Metabolite standards were purchased from Sigma-Merck (Singapore). Amino acids and stable-isotope internal standard mixes (ISTDs) were purchased from Cambridge Isotopes laboratories (Massachusetts, USA). Organic solvents of LC–MS grade were purchased from Aik Moh Chemicals Pte Ltd. (Singapore). Milli-Q IQ7000 LC-MS grade water and PEEK-lined SeQuant^®^ZIC^®^-cHILIC 3um,100Å 100 x 2.1 mm HPLC column were purchased from Merck Pte Ltd, Singapore.

Oocytes, calibrators, and quality controls were placed in PCR tubes, supplemented with 10 µL of a 5 mg/L heavy isotope internal standards mixture. Following this, 50 µL of ice-cold acetonitrile containing 0.1% formic acid was added to each tube, mixed at 1000 rpm/min for 10 minutes using a shaker, and transferred to Eppendorf tubes. Samples were then centrifuged at 13,800 xg for 10 minutes at 4°C. The supernatant was carefully transferred to a 96-microwell plate and loaded into the auto-sampler (8 °C), and 10 µL was used for LC-MS/MS analysis. Ion counts from the peak area under the curve (AUC) were normalized against the heavy isotope internal standards. The comprehensive analysis of small metabolites in the oocytes was performed using the Agilent 1290 Infinity II/Agilent 6495A Triple Quadrupole LC/MS system operated with the electrospray ionization (ESI) in either positive or negative ionization mode as reported previously ^68,69^. Acquisitions, qualitative and quantitative analysis were performed using the Agilent MassHunter Workstation Acquisition (10.0.127), Agilent Qualitative Analysis (10.0), and Agilent Quantitative Analysis (10.0) software. To ensure high-quality data for statistical analysis and interpretation for all metabolites, the following steps and measures were taken: (1) the samples were analysed within a single batch, (2) the injection order of samples was randomized, (3) carry-over was checked by injecting water blanks between samples, (4) background noises were excluded from area integration, (5) peak picking was by m/z ratio and retention time (RT), and (6) RT correction and m/z peak alignment across multiple spectra were performed.

All statistical analysis for MS experiments were performed with R (4.2.2). Area response ratios were log2-transformed and fitted to generalized linear models for group-wise comparison with function glm. Only metabolites reaching statistical significance (p-value of Wald test < 0.05) in this univariate analysis were included in subsequent analysis. Partial least squares-discriminant analysis (PLS-DA) was performed with the “mixOmics” and “ropls” packages. To assess model fit and avoid overfitting, the predictive performance of the model was assessed with leave-one-out cross-validation (LOOCV), and significance was validated with permutation tests (p-value < 0.001 for R^2^X_cum_ and Q^2^_cum_). The three-dimensional score plot of the fitted PLS model was visualized with Asymptote vector graphics language (https://asymptote.sourceforge.io/). Heatmap was constructed with the “pheatmap” package with reference to z-score in each metabolite. Package “tidyverse” was used for data wrangling purposes.

### Measurement of oocyte ATP levels

Total ATP content in a pool of 5 oocytes was determined using the Adenosine 5′-triphosphate (ATP) bioluminescent somatic cell assay kit (FLASC, Sigma) following the manufacturer’s instructions. Briefly, 100 μl of ATP Assay Mix Working Solution (1:25 diluted from ATP Assay Mix Stock Solution) was added to a 96-well plate (reaction vial) and allowed to stand at room temperature for 3 minutes. Somatic Cell ATP Releasing Reagent (100 μl), 50 μl filtered ultrapure water and 50 μl sample were added to a new tube and 100 μl was transferred to the reaction vial. ATP concentration was measured immediately using plate reader (ThermoFisher Varioskan LUX).

### Image analysis and quantification

Images were analyzed using ImageJ (NIH), Imaris (Bitplane), or ZEN 3.2 (Zeiss) software. The exported data were further processed in Excel, GraphPad Prism 9, or Origin 2021b.

To measure the follicle diameter, the line tool in ImageJ was utilized. Firstly, a line was drawn across the widest diameter of the follicle, and the length of the line was noted. Subsequently, a second line was drawn across the diameter of the follicle, perpendicular to the first line, and the length of the line was noted. The final follicle diameter was determined by calculating the average length of these two measurements.

To quantify Rec8 cohesin intensities within chromosomes, z-stack images of chromosome (stained with DAPI) and rec8 staining were processed and analyzed using Imaris. First, the chromosomes were identified and delineated using the “Surface” function tailored to the chromosome channel. This generated surface was then utilized to isolate the Rec8 signal, employing the “mask all” function, effectively segregating the Rec8 signal present within the chromosome confines. The isolated Rec8 signal was then channeled separately. Using the “Spatial - Intensity inside” function specific to the chromosome surface, the subsequent average intensity of this newly defined Rec8 channel was computed.

Sister kinetochore distance analyses for metaphase I and metaphase II oocytes were conducted on super resolution images of 2mEGFP-CENP-C and H2B-mCherry obtained using LSM980 microscope equipped with an AiryScan2 detector. To identify sister kinetochores at the metaphase I stage, images were subjected to deconvolution using Huygens Professional software (Scientific Volume Imaging b.v.) to improve resolution. The microscopy type was specified as “Array Detector Confocal (Zeiss Airyscan2)”, and metadata were imported from image files. A theoretical PSF was calculated using the provided information, followed by the application of the iterative maximum likelihood estimation CMLE algorithm for deconvolution. Deconvolved images were subsequently analyzed using Imaris (Bitplane) with the “Spots” function. For kinetochore identification at the metaphase I stage, XY and Z diameters were set to 0.125 µm and 0.375 µm, respectively. Spots were chosen based on the quality criterion algorithm, and the distance to the nearest neighbor were calculated and assessed.

For sister kinetochore identification at the MII stage, AiryScan-processed images were acquired using ZEN blue software and were subsequently analyzed using Imaris with the “Spots” function, where XY and Z diameters were adjusted to 0.3 µm and 0.5 µm, respectively. Spots were again selected using the quality criterion algorithm, and sister kinetochore distances were ascertained via the Measurement Points function. The line mode was designated as “pairs,” with paired spots manually selected. Center-to-center distances between paired spots were subsequently determined and analyzed. PSSC was excluded from the distance measurement process.

To measure the fluorescence intensity of JC-1, TMRM, CM-H2DCFDA, and MitoSOX Red in live oocytes, ImageJ software was employed. An area was designated as the region of interest (ROI) to include the entire oocyte. The mean fluorescence intensity for each unit area within this specified ROI was then determined.

To quantify spindle abnormalities, we classified spindles exhibiting elongation, shortening, collapse, multipolar or monopolar, and unfocused spindle poles as abnormal, consistent with criteria established in previous studies^63,70–72^.

To quantify the density of TZPs, airyscan super-resolution images of phalloidin-stained TZPs were analyzed using ImageJ, focusing on a confocal optical section taken at the oocyte’s equatorial plane. Utilizing four 10-μm arcs spaced 90° apart around the oocyte’s circumference, the number of TZPs was then manually counted within each of these 10-μm arcs. To determine the average density of TZPs, the total counts from all four arcs were summed and then divided by four, providing the mean number of TZPs per 10 μm segment.

### Statistical analyses

Statistics were calculated using GraphPad Prism 9 software or R software. Statistical significance for two-group comparisons was determined using two-tailed Student’s t-tests or two-tailed Fisher’s exact test; for three-group comparisons, one-way analysis of variance (ANOVA) with Tukey’s multiple comparison test was used, unless indicated otherwise in the figure legends or methods section. For the boxplots in the figures, black square and centre bars indicate mean and median of the population, respectively. The upper and lower hinges represent the third and the first quartiles; the whiskers show a 1.5x interquartile range. For the violin plot, the shaded area represents the data distribution. Box plots inside the violins indicate mean and median by black circle and center bar; upper and lower limits and whiskers represent quartiles and 1.5 × interquartile. Sample sizes or numbers of independent experiments are indicated in figures or figure legends. All tests were performed with a significance level of *P* < 0.05. Data presented are derived from at least three independent experiments. In the figures, significance is designated as follows: \*\*\*\**P* < 0.0001; \*\*\**P* < 0.001; \*\**P* < 0.01; \**P* <0.05; ns, *P* > 0.05.

## Data availability

All data needed to evaluate the conclusions in the study are present in the main text or the supplementary materials.

---

**Extended Data Figure 1.**
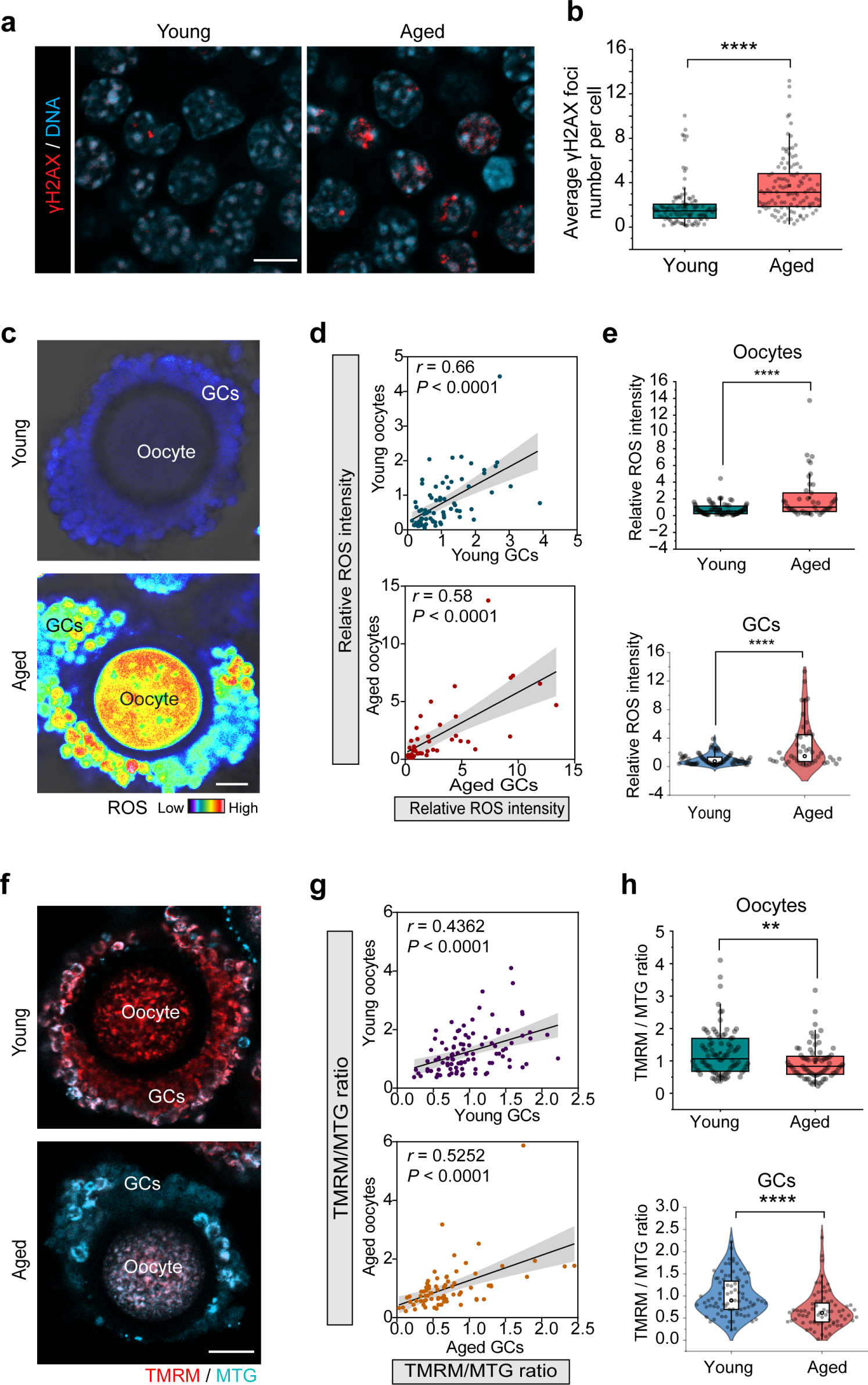
Follicles accumulate age-related abnormalities. **a.** Representative images of γH2AX staining (red) in granulosa cells from young and aged follicles. DNA stained with DAPI. Scale bar, 10 μm. **b.** Quantification of γH2AX foci in granulosa cells from young (2 month) and aged (14 month) follicles. **c.** CM-H2DCFDA staining to visualize ROS levels in *in vivo* isolated young and aged oocyte-GC complexes. Scale bar, 30 μm. **d.** Scatter plot demonstrating the correlation between ROS levels in granulosa cells and oocytes, with the gray area indicating the 95% confidence intervals. The Pearson’s correlation coefficient (r) is displayed. n = 76 (young), 43 (aged). **e.** Fluorescence intensity of ROS signals in oocytes (upper panel) and GCs (lower panel), respectively. 2-month-old (young) and 14-month-old (aged) wide-type ICR mice were used. **f.** Fluorescence images of young and aged *in vivo* isolated oocyte-GC complexes stained with MitoTracker Green (MTG, cyan) and mitochondrial membrane potential-sensitive dye TMRM (red). n = 84 (young), 72 (aged). Scale bar, 30 μm. **g.** Scatter plot showing the correlation between mitochondrial membrane potential (ΔΨm) in granulosa cells and oocytes, with the grey area indicating the 95% confidence intervals. ΔΨm was determined as the mean fluorescence intensity ratio of TMRM to MTG. The Pearson’s correlation coefficient (r) is shown. **h.** Quantification of the fluorescence intensity ratio of TMRM to MTG in oocytes and granulosa cells. 2-month-old (young) and 14-month-old (aged) wide-type ICR mice were used. Box plots, mean and median are indicated by black square and centre bar; upper and lower limits and whiskers are quartiles and 1.5 × interquartile range. Two-tailed unpaired t-tests for (**b**), (**e**), and (**f**). \*\*\*\**P* < 0.0001; \*\**P* < 0.01.

**Extended Data Figure 2.**
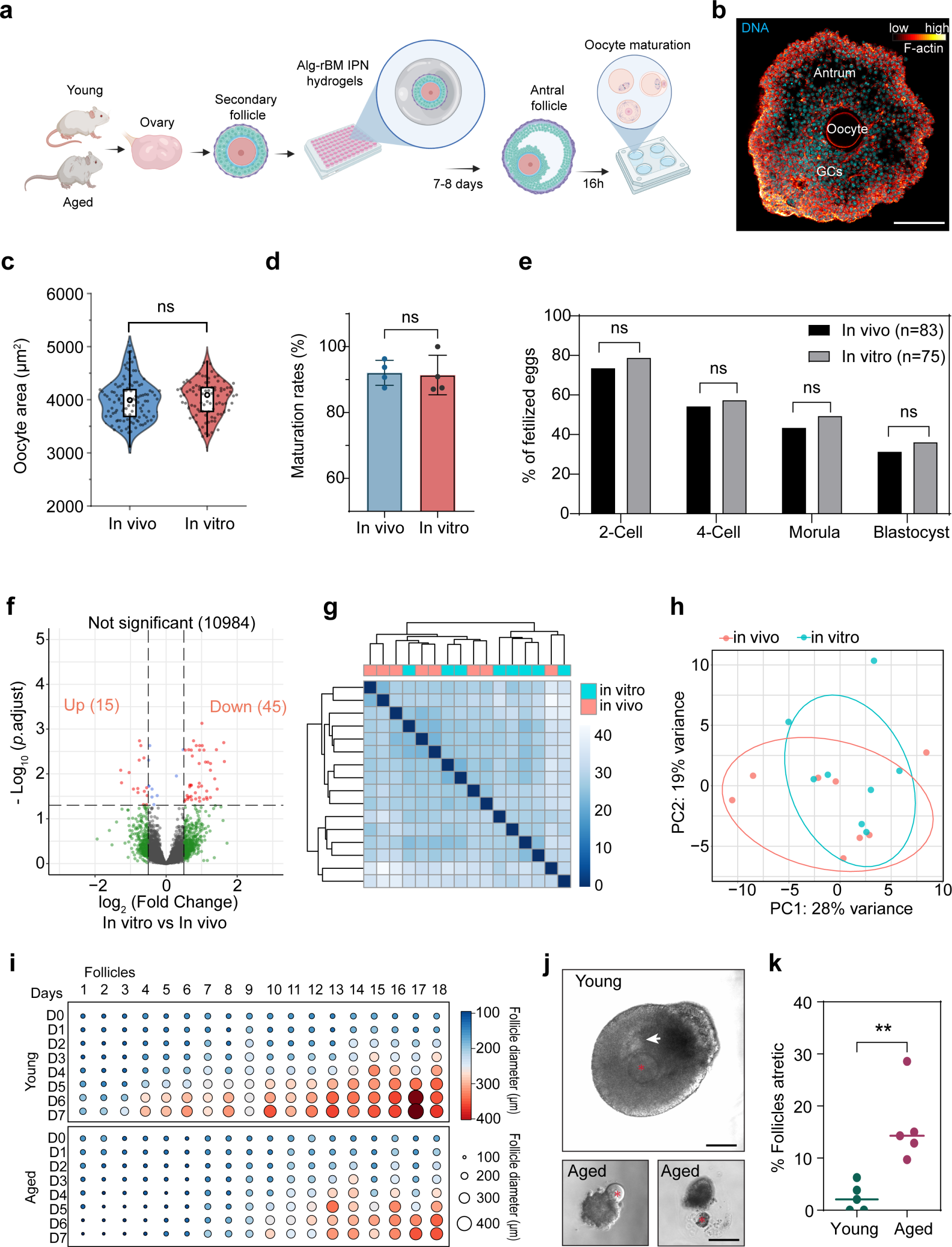
Comparison of *in vivo* and *in vitro* grown oocytes. **a.** Schematic representation of the 3D *ex vivo* follicle culture system using Alg-rBM IPN hydrogels (See Methods for details). **b.** Representative immunofluorescence image of a 3D *ex vivo* cultured antral stage follicle. Follicle was stained with phalloidin to label F-actin, DNA was stained with DAPI. Scale bar, 100 μm. **c.** Diameter of oocytes grown *in vivo* or *in vitro*. n = 109 (*in vivo*), 90 (*in vitro*). 2-3 month-old wide type ICR mice were used. Two-tailed unpaired t-tests. ns, *P* > 0.05. **d.** Quantification of oocyte maturation rate (PB1 extrusion rate) among *in vitro* and *in vivo* oocytes. n = 126 (*in vivo*), 98 (*in vitro*). 2-3 month-old wide type ICR mice were used. Two-tailed unpaired t-tests. ns, *P* > 0.05. **e.** Analysis of embryo development potential following *in vitro* fertilization (IVF) of oocytes grown *in vivo* or *in vitro*. n = 83 (*in vivo*) and 75 (*in vitro*). 2-3 month-old wide type ICR mice were used. Fisher’s exact test. ns, *P* > 0.05. **f.** Transcriptome analysis of oocytes grown *in vitro* and *in vivo*. Differentially expressed genes (p.adjust < 0.05 and log_2_ fold change > 0.5 or < −0.5) in oocytes cultured *in vitro* based on evaluation using the DESeq2 algorithm. Only about 0.5% (60 genes out of 11044) of genes were differentially expressed between oocytes grown *in vitro* and *in vivo*. For the transcriptome analysis, 6 *in vivo* and 8 *in vitro* fully grown germinal vesicle (GV) oocytes were used. **g.** Correlation heatmap with hierarchical clustering to show the sample-to-sample distances calculated from transcriptome data of *in vitro* and *in vivo* grown oocytes. **h.** Principal component analysis (PCA) of the normalized gene expression data from *in vivo* and *in vitro* grown oocytes. Each dot represents a single oocyte from each group. Ellipses fit a multivariate t-distribution at confidence level of 0.8. **i.** Dot plots illustrating changes in the size of young and aged follicles over time during 3D *ex vivo* culture. The color bar and circle size represent the follicle size. The diameters of 18 young and aged follicles were monitored daily from day 0 to day 7. 2-month-old (young) and 14-month-old (aged) wide-type ICR mice were used. **j.** Follicles from aged mice were more prone to undergo atresia during *in vitro* culture. Follicles from young mice (top) maintained their 3D structure with proliferation of GCs and antrum formation (white arrowhead) while being cultured in Alg-rBM IPN. In contrast, follicles from aged mice experienced increased atresia during *in vitro* culture (bottom). Follicles were considered atretic if there was disruption of contact between the oocyte (red asterisk) and granulosa cells, leading to the release of oocytes from the follicles (bottom left), or if the follicles contained apoptotic or dead oocytes (bottom right). Scale bar, 100 μm. **k.** Atresia rate was quantified in young (2-3 month) and aged (14-15 month) follicles after 3D *in vitro* culture. The median is represented by the center line. n = 166 (young), 199 (aged). Two-tailed unpaired t-tests. \*\**P* < 0.01.

**Extended Data Figure 3.**
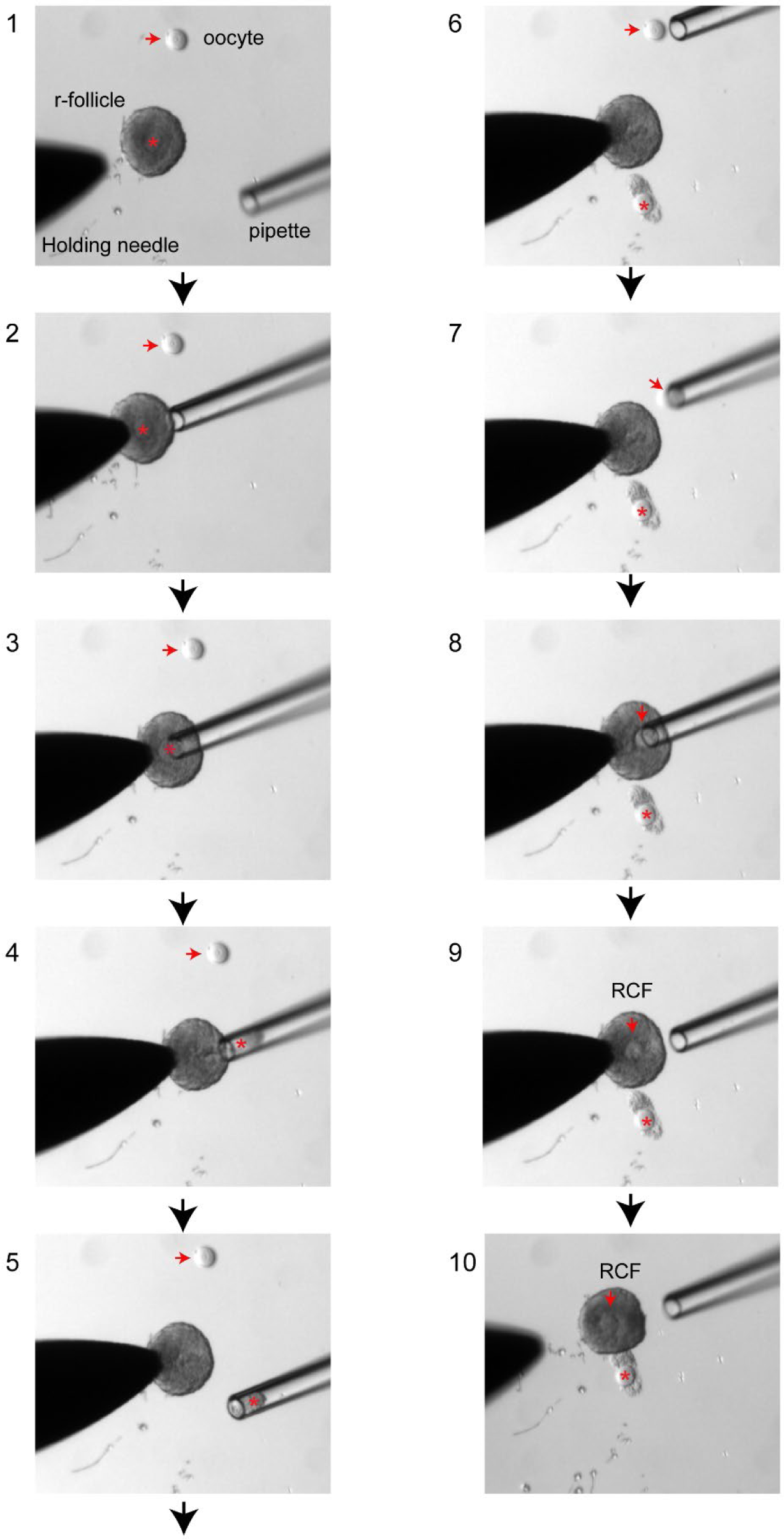
Procedure for generating reconstituted chimeric follicles. Red arrow points to the oocyte used for transplantation. Red asterisk indicates the oocyte within the r-follicle that will be replaced. Refer to Supplementary Video 1 and Methods for further details.

**Extended Data Figure 4.**
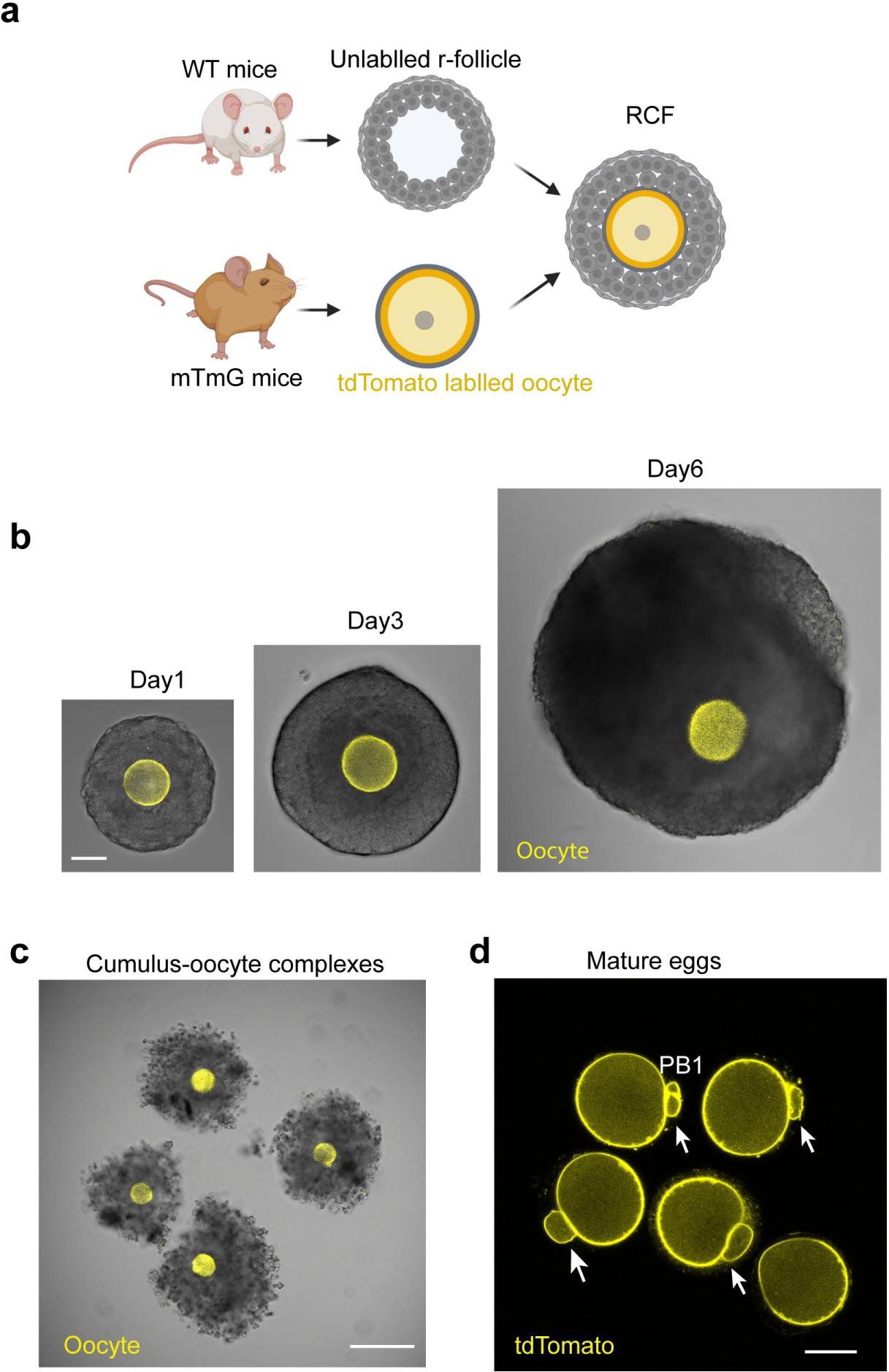
Growth and maturation of RCFs in 3D *ex vivo* culture. **a.** To distinguish between the donor oocyte and the r-follicle, we employed oocytes from mTmG transgenic mice exhibiting membrane-localized tdTomato (pseudo-colored yellow). In contrast, the r-follicles were sourced from non-fluorescent wild-type mice. The mTmG oocytes served as donors as referenced in Fig. 1i and Extended Data Fig. 4. **b.** RCF size increase during 3D ex vivo culture. Oocytes from transgenic mTmG mice and follicular somatic cells from wild-type mice, as shown in (**a**). Scale bars, 50 μm. **c.** Cumulus-oocyte complexes (COCs) isolated from antral RCFs were induced for oocyte maturation with hCG for 16 hours *in vitro*. Note that cumulus cells surrounding the oocytes (from mTmG mice) expanded, and oocytes resumed meiosis, extruded the PB1 as shown in (**d**). Scale bars, 200 μm. **d.** Representative image of mature eggs derived from RCFs as shown in (**b** and **c**). The cumulus cells were removed after maturation to visualize mature eggs with the first polar body (PB1, arrows). Scale bars, 40 μm.

**Extended Data Figure 5.**
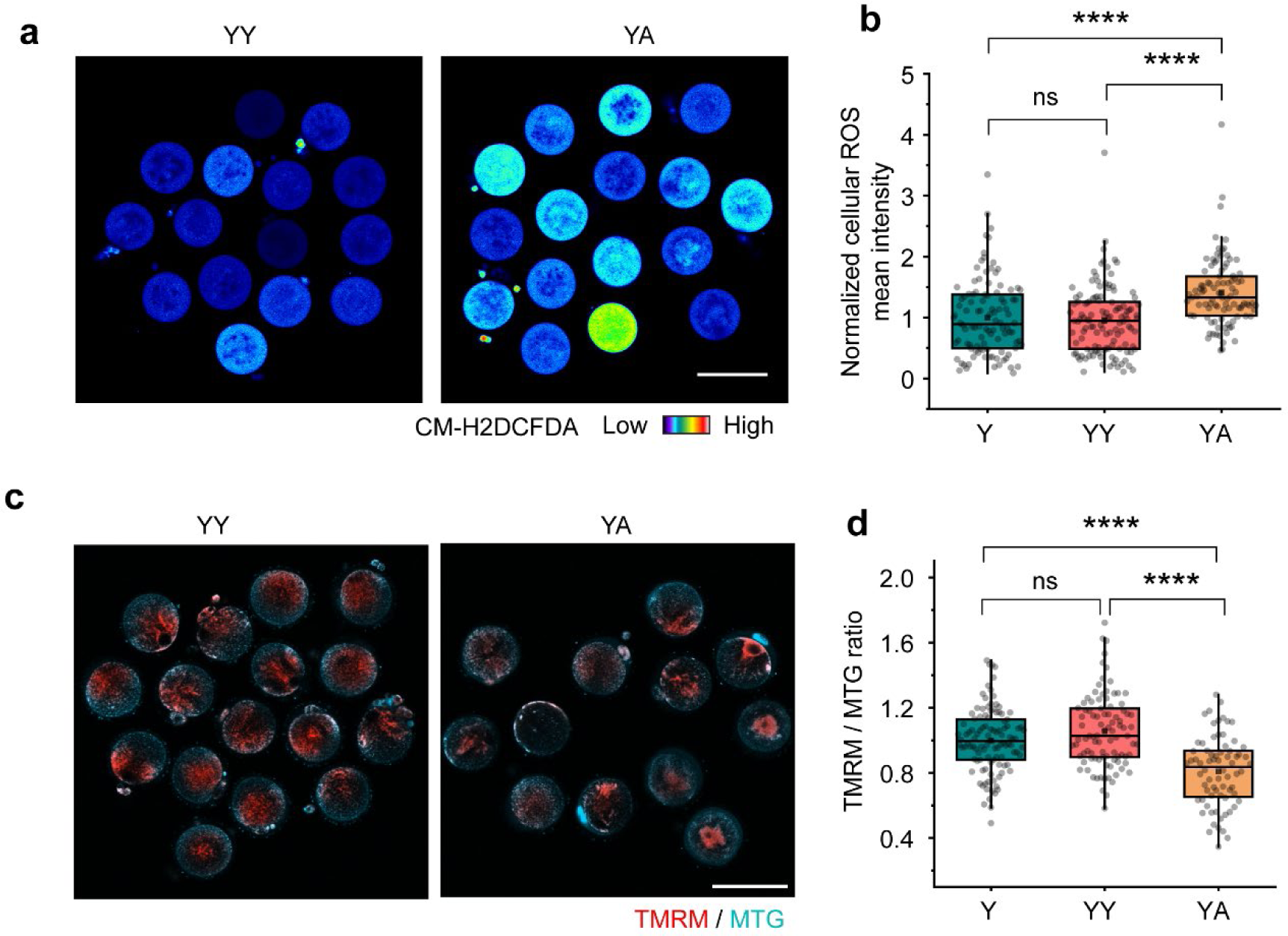
Aged follicular somatic cells elevate ROS levels and reduce mitochondrial membrane potential in young oocytes. **a.** Representative confocal images of cellular ROS stained with CM-H2DCFDA in oocytes from YY and YA RCFs. Scale bar, 100 μm. **b.** Quantification of CM-H2DCFDA fluorescence intensity in oocytes from YY and YA RCFs, as well as Y. n = 105 (Y), 122 (YY), 97 (YA). 2-month-old (young) and 14-month-old (aged) wide type ICR mice were used. **c.** Fluorescence images of oocyte stained with MitoTracker Green (MTG, cyan) and mitochondrial membrane potential-sensitive dye TMRM (red). Scale bar, 100 μm. **d.** Quantification of the fluorescence intensity ratio of TMRM to MTG in oocytes from YY and YA RCFs, as well as Y. n = 110 (Y), 94 (YY), 72 (YA). 2-month-old (young) and 14-month-old (aged) wide type ICR mice were used. Box plots, mean and median are indicated by black square and centre bar; upper and lower limits and whiskers are quartiles and 1.5 × interquartile range. One-way ANOVA, Tukey’s multiple comparisons test for (**b**) and (**d**). \*\*\*\**P* < 0.0001; ns, *P* > 0.05.

**Extended Data Figure 6.**
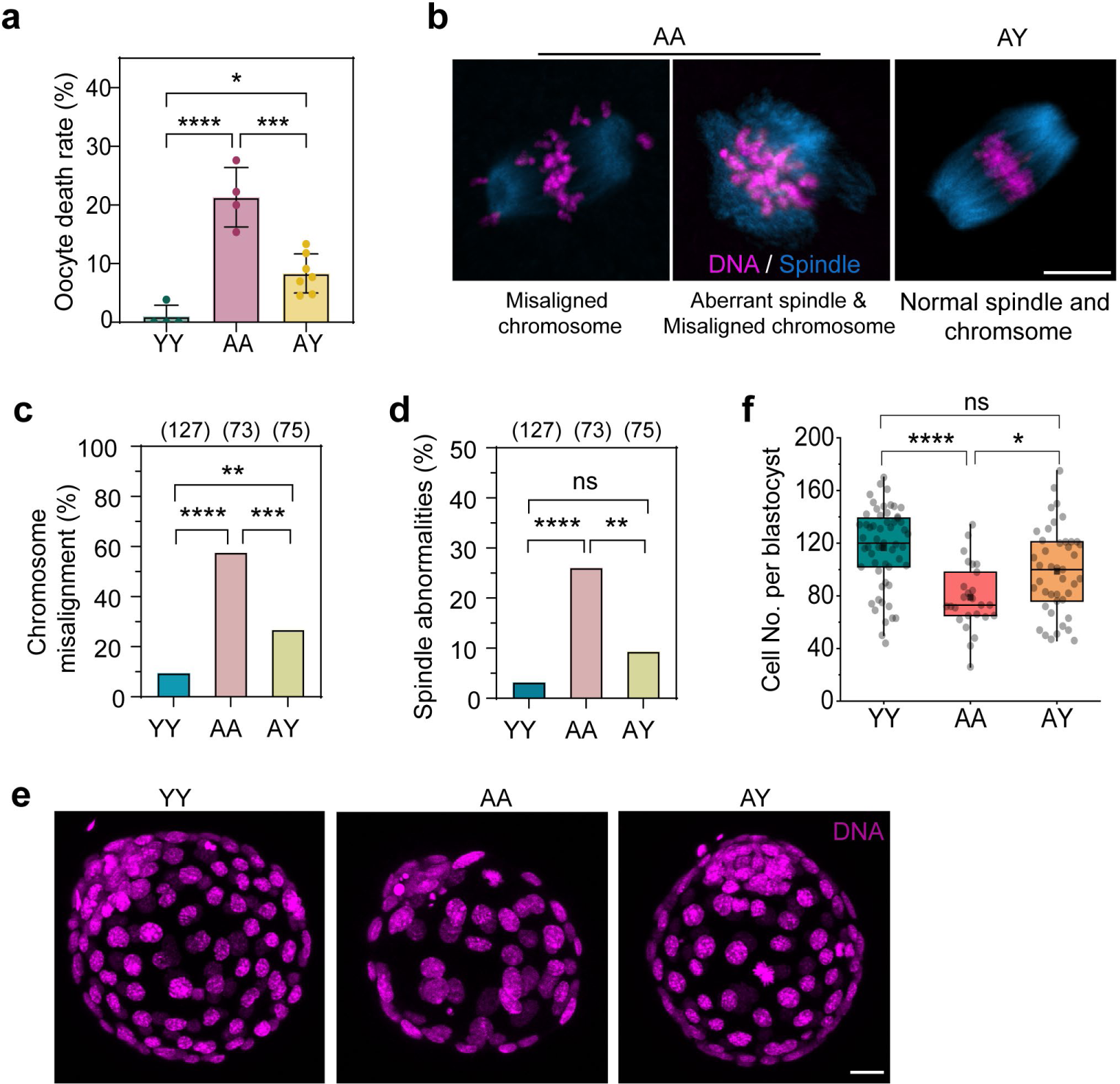
Impact of young follicular somatic cells on aged oocyte chromosomal stability and blastocyst cell numbers. **a.** Oocyte death rates were quantified in oocytes from YY, AA, and AY RCFs. Data are shown as mean ± SD. n = 77 (YY), 142 (AY), 65 (AA). 2-3 month-old (young) and 14-15 month-old (aged) wide type ICR mice were used. **b.** Representative live cell images of meiotic spindle and chromosomes in MII oocytes from AA and AY RCFs. The left image shows an example of chromosomal misalignment in aged oocyte from AA RCF, the image in middle shows an example of abnormal spindle in an aged oocyte from an AA RCF, while the right image shows example of a normal spindle in aged oocyte from AY RCF. Spindles were stained with SiR-tubulin, and chromosomes (DNA) were stained with Hoechst. Scale bar, 10 µm. **c,d.** Quantification of the percentage of chromosomal misalignment (**c**) on the metaphase plate and spindle abnormalities (**d**) in oocytes from YY, AA and AY RCFs. The numbers of oocytes are specified in brackets. 2-3 month-old (young) and 14-17 month-old (aged) wide type ICR mice were used. Fisher’s exact test. **e.** Representative images of DAPI-stained blastocysts for cell number counting. Scale bars, 20 μm. **f.** Cell numbers per blastocyst were quantified. n = 57 (YY), 43 (AY), 26 (AA). Box plots depict data distribution, where the mean and median are indicated by black square and center bar; upper and lower limits and whiskers are quartiles and 1.5 × interquartile range. One-way ANOVA, Tukey’s multiple comparisons test for (**a**) and (**f**). \*\*\*\**P* < 0.0001; \*\*\**P* < 0.001; \*\**P* < 0.01; \**P* <0.05; ns, *P* > 0.05.

**Extended Data Figure 7.**
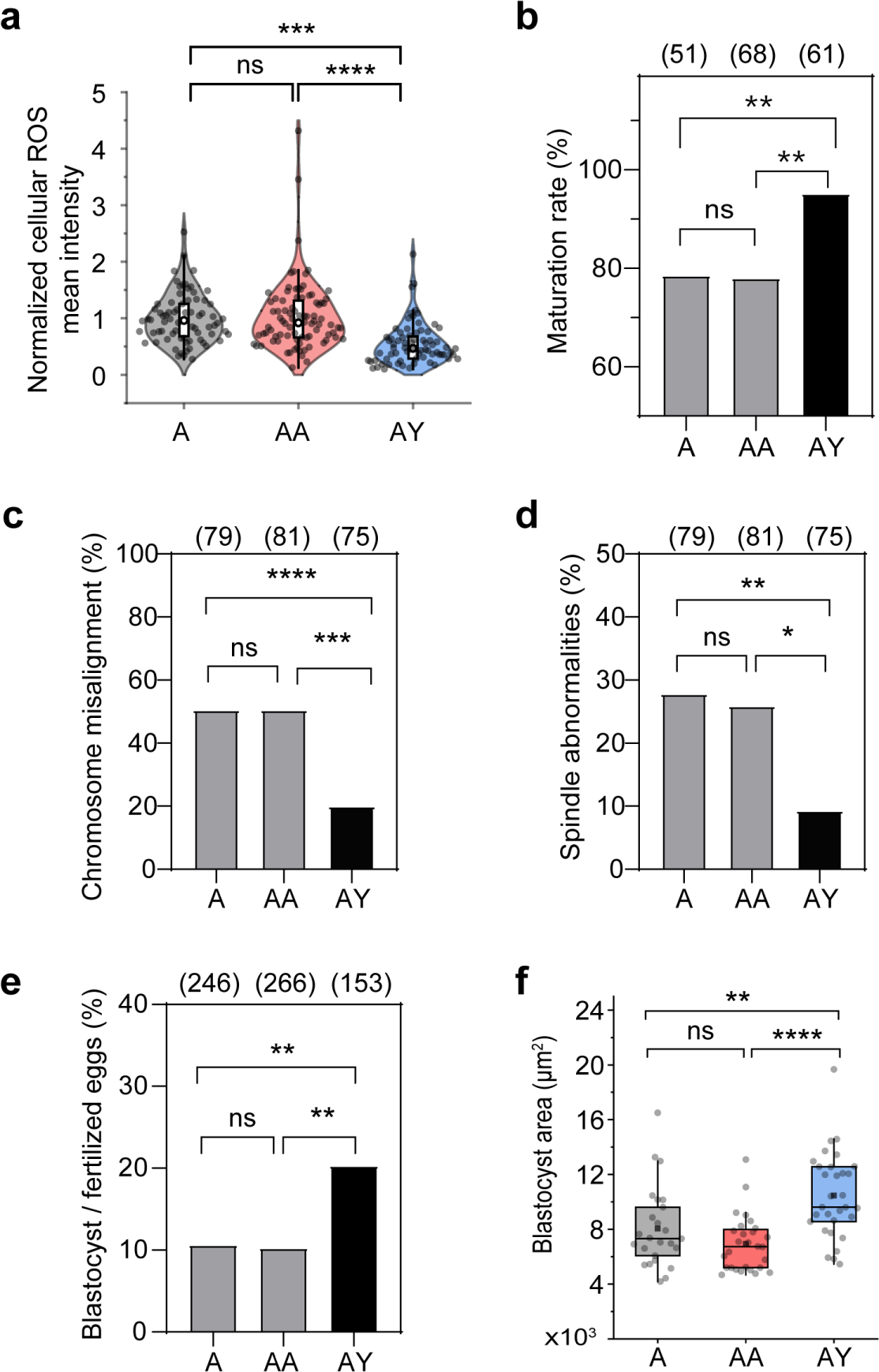
Comparative analysis of oocytes from AA RCFs and aged intact follicles (A). **a.** Quantification of cellular ROS levels in oocytes from AA and AY RCFs as well as from aged intact follicles (A). n = 72 (A), 72 (AA), 72 (AY). Cellular ROS was stained with CM-H2DCFDA. 2-month-old (young) and 14-month-old (aged) wide type ICR mice were used. One-way ANOVA, Tukey’s multiple comparisons test. **b.** Comparison of oocyte maturation rates (PB1 extrusion rates) among oocytes from AA and AY RCFs as well as from aged intact follicles (A). The number of oocytes with the PB1 extruded was counted after maturation for 16h. 2-month-old (young) and 14-month-old (aged) wide type ICR mice were used. The numbers of oocytes are specified in brackets. Fisher’s exact test. **c,d.** Quantification of the percentage of chromosomal misalignment (**c**) on the metaphase plate and spindle abnormalities (**d**) in oocytes from AA and AY RCFs as well as from aged intact follicles (A). The numbers of oocytes are specified in brackets. 2-month-old (young) and 14-month-old (aged) wide type ICR mice were used. Fisher’s exact test. **e.** Analysis of blastocyst formation rate following *in vitro* fertilization (IVF) of oocytes from AA and AY RCFs as well as from aged intact follicles (A). n = 246 (A), 266 (AA), 153 (AY). Fisher’s exact test. 2-month-old (young) and 14-month-old (aged) wide type ICR mice were used. **f.** Quantification of blastocyst size measured as projected area. n = 25 (A), 28 (AA), 31 (AY). Box plots indicate mean and median by black square and center bar; upper and lower limits and whiskers represent quartiles and 1.5 × interquartile; one-way ANOVA, Tukey’s multiple comparisons test. 2-month-old (young) and 14-month-old (aged) wide type ICR mice were used. \*\*\*\**P* < 0.0001; \*\*\**P* < 0.001; \*\**P* < 0.01; \**P* <0.05; ns, *P* > 0.05.

**Extended Data Figure 8.**
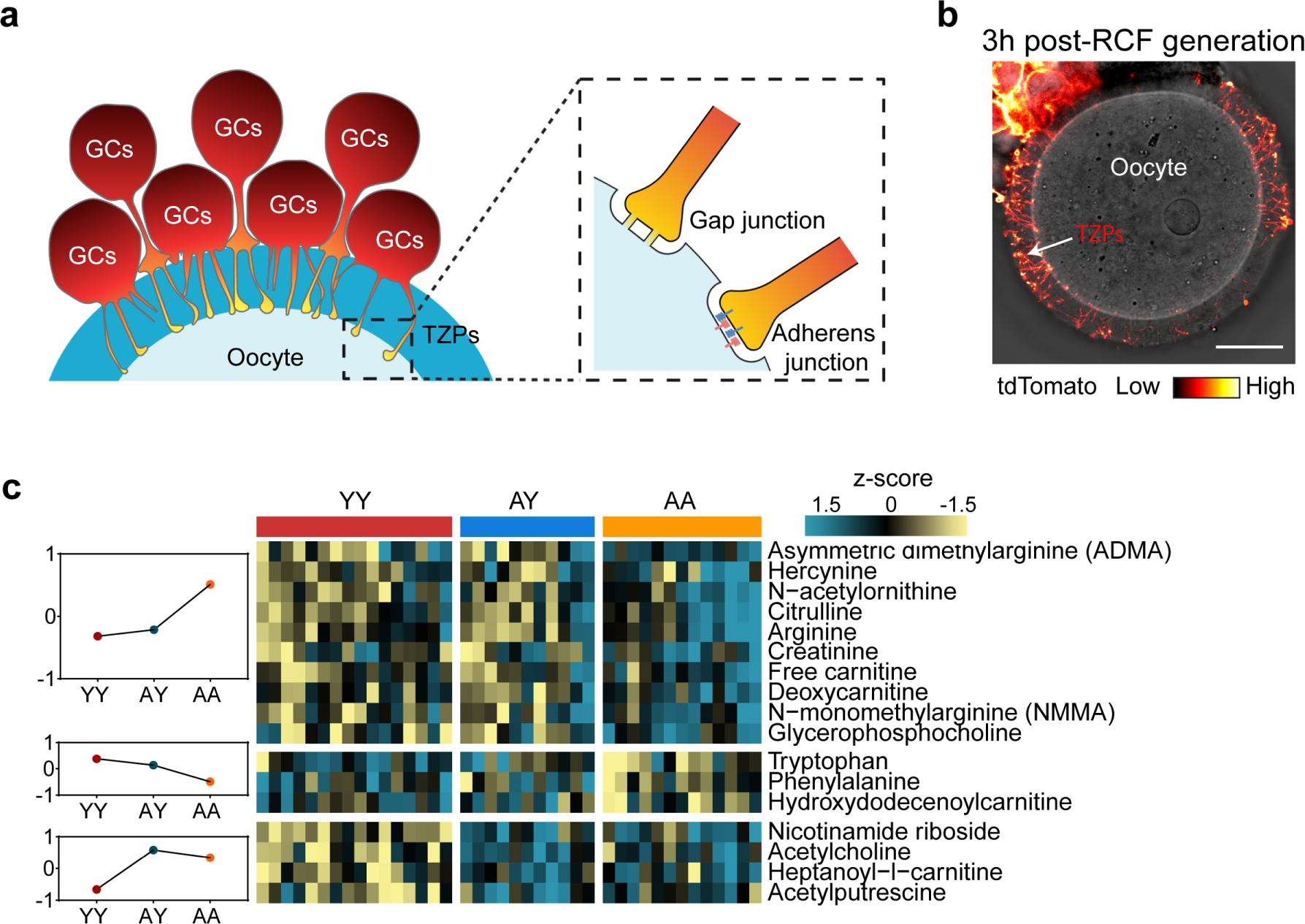
TZP regeneration and metabolic remodeling of oocytes in RCFs. **a.** Schematic demonstrating TZPs from GCs that pass through the zona pellucida, forming either adherens junctions or gap junctions on the oocyte surface. **b.** TZP regenerated within 3 hours of RCF culturing. RCF containing follicular somatic cells from mTmG mouse and wild-type oocytes were cultured within Alginate-rBM beads for 3h. Somatic cells were then removed to visualize TZP regeneration. Scale bars, 20 μm. **c.** Heatmap and line plot of differential metabolites in oocytes from YY, AA, and AY RCFs. Heatmap displayed the 17 differential metabolites (*P* < 0.05) between YY and AA oocytes, with 13 of them being rescued in AY oocytes (those in the top two blocks). Group-wise differences were summarized with line plot on the left.

**Extended Data Figure 9.**
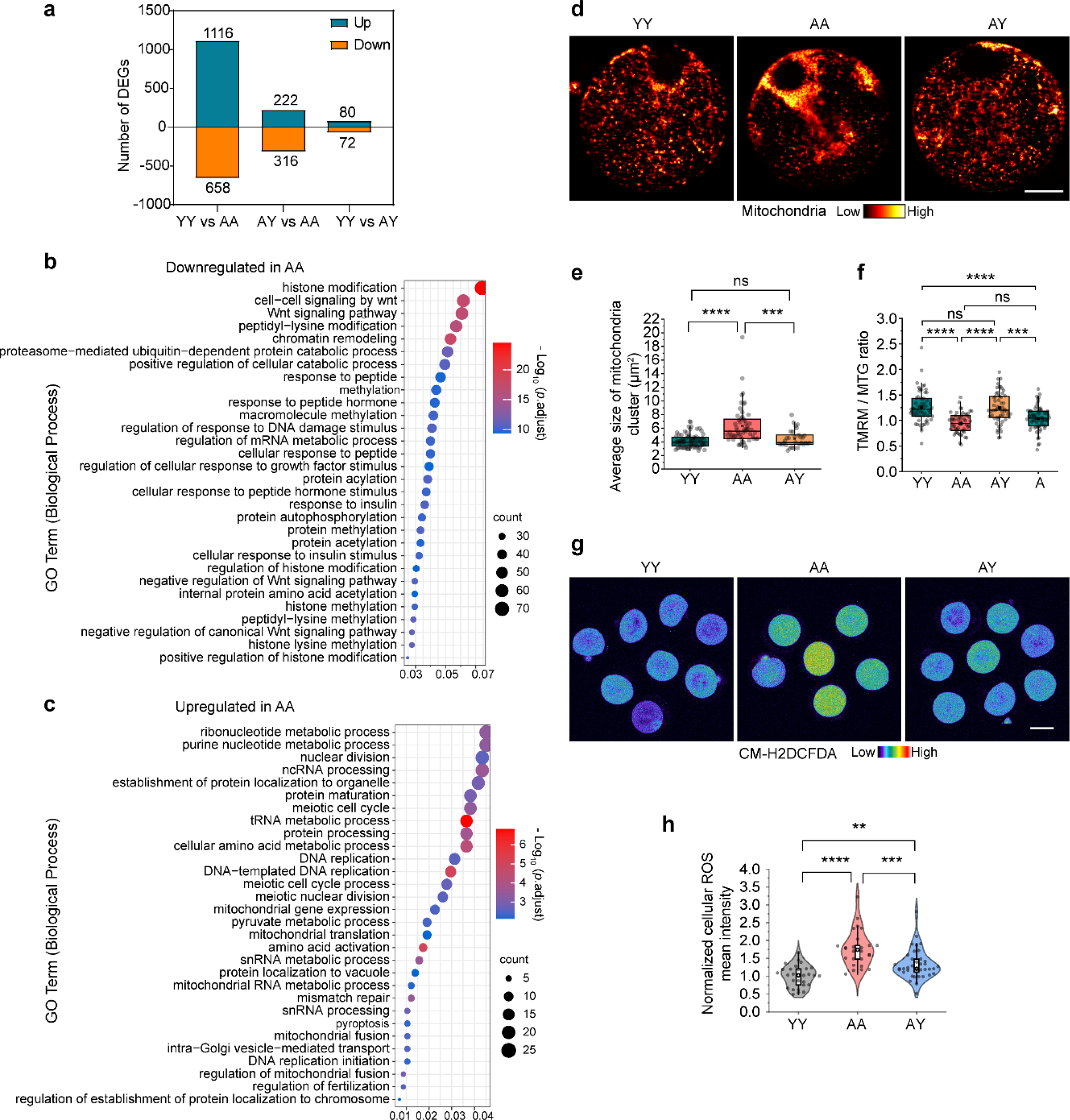
Young somatic cells restore mitochondrial fitness in aged oocytes. **a.** Histogram displays the number of up-regulated or down-regulated DEGs between oocytes from YY and AA, AY and AA, or YY and AY RCFs. **b,c.** Representative GO terms associated with the genes that were downregulated (panel b) and upregulated (panel c) in aged oocytes from AA RCFs when compared to young oocytes from YY RCFs **d.** Confocal microscopy images of mitochondrial distribution patterns in oocytes from YY, AA, and AY RCFs. Mitochondria were detected by using MitoTracker Red CMXRos. Scale bar, 20 μm. **e.** Mitochondria cluster sizes in oocytes from YY, AA, and AY RCFs were calculated and compared. n = 73 (YY), 55 (AA), 31 (AY). 2-month-old (young) and 14-15 month-old (aged) wide type ICR mice were used. Box plot displays data distribution, with the mean and median represented by a black square and center bar, respectively; the upper and lower limits, as well as whiskers, correspond to quartiles and 1.5 × the interquartile range. Scale bar, 20 μm. **f.** Quantification of the mitochondria membrane potential using fluorescence intensity ratio of TMRM to MTG (MitoTracker Green). n = 59 (YY), 53 (AA), 57 (AY), 61 (A). 2 month-old (young) and 14 month-old (aged) wide type ICR mice were used. **g.** Representative confocal images of cellular ROS stained with CM-H2DCFDA in oocytes from YY, AA, and AY RCFs. Scale bar, 50 μm. **h.** Quantification of CM-H2DCFDA fluorescence intensity in oocytes from YY, AA, and AY RCFs. n = 28 (YY), 26 (AA), 39 (AY). 2-month-old (young) and 14-15 month-old (aged) wide type ICR mice were used. One-way ANOVA, Tukey’s multiple comparisons test for (**e**), (**f**), and (**h**). \*\*\*\**P* < 0.0001; \*\*\**P* < 0.001; \*\**P* < 0.01; ns, *P* > 0.05.

**Extended Data Figure 10.**
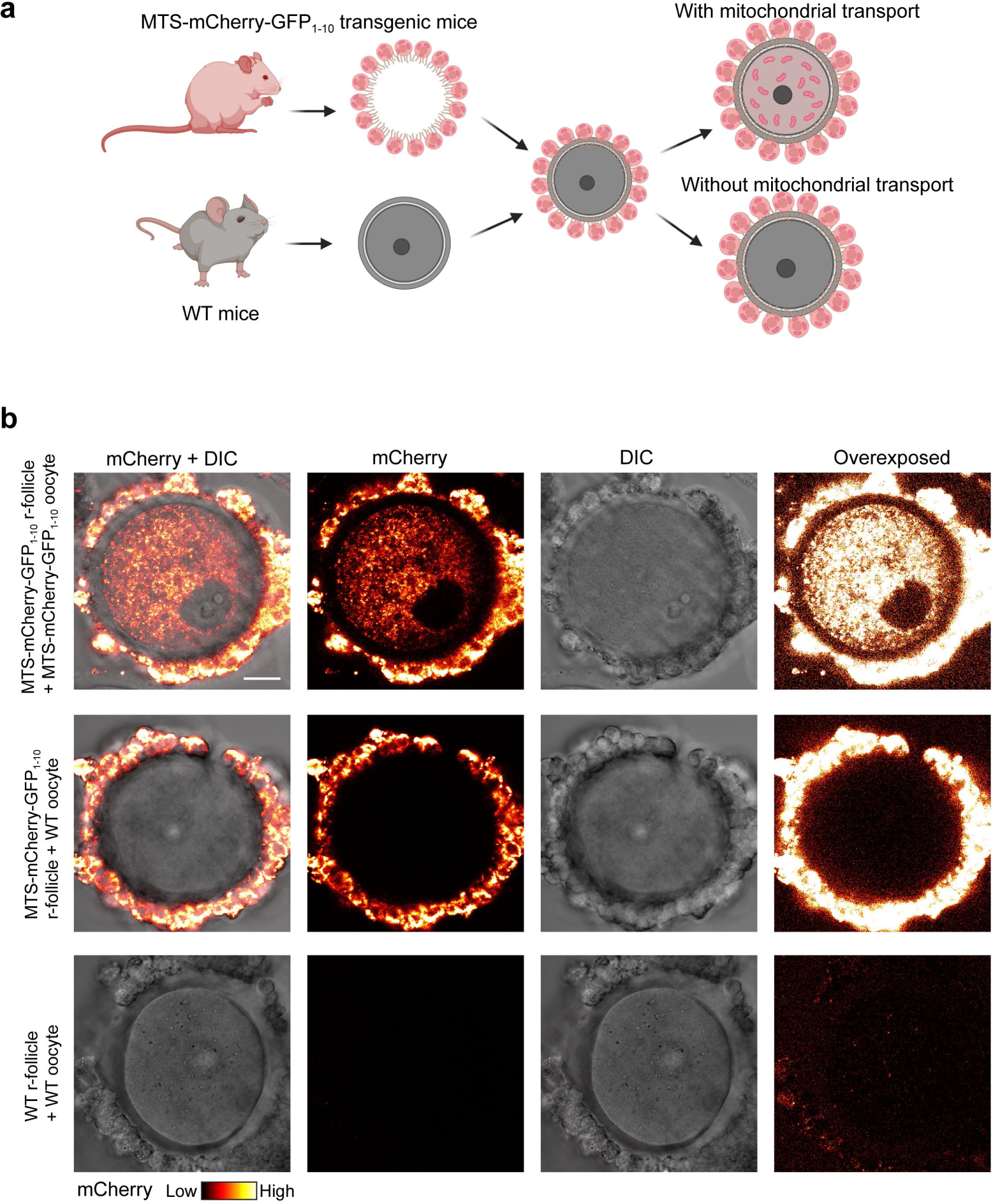
Investigating possible GC-to-oocyte mitochondrial transport in RCFs using MTS-mCherry-GFP_1-10_ transgenic mice. **a.** Experimental design to study mitochondrial transport within RCFs. RCFs were created using somatic cells from transgenic MTS-mCherry-GFP_1-11_ mice, which express mitochondria-targeted mCherry, and unlabelled oocytes from wild-type mice. **b.** For a positive control, confocal microscopy images of mCherry-labelled mitochondria in oocytes. Top panel: An RCF formed by transplanting an MTS-mCherry-GFP_1-11_ oocyte into an MTS-mCherry-GFP_1-10_ r-follicle. Middle panel: An RCF generated by transplanting a wild-type oocyte into an MTS-mCherry-GFP_1-10_ r-follicle. Bottom panel: For a negative control, an RCF generated by transplanting a wild-type oocyte into a wild-type r-follicle, serving as a negative control. Right most panel of each row: overexposed images corresponding to images of the second column (mCherry). Somatic cells were partially removed before imaging to better observe oocyte fluorescence. Scale bar, 20µm.

**Extended Data Figure 11.**
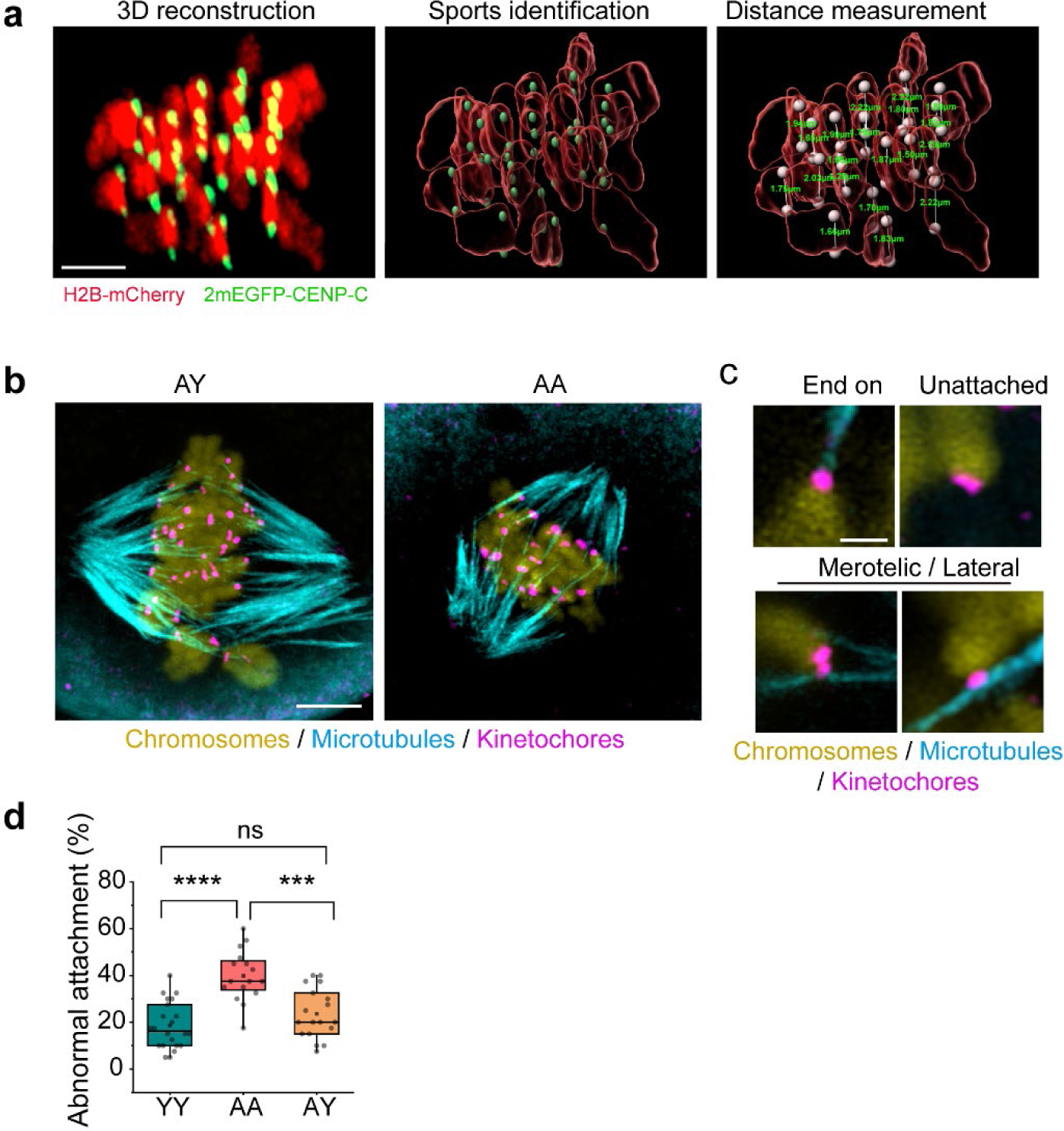
Reduced kinetochore-microtubule attachment defects in aged oocytes from AY RCFs. **a.** Representative example of measurements of sister kinetochore pair distance in MII oocytes expressing 2mEGFP-CENP-C and H2B-mCherry to label kinetochores and chromosomes, respectively. Images were generated by using Imaris software (see Methods for details). Also see Supplementary Video 2. Scale bar, 2 µm. **b.** Representative confocal images of cold shock-treated oocytes 5.5h after GVBD. Images are projections of a confocal z-series showing microtubules (alpha-tubulin, cyan), kinetochores (CREST, magenta), and DNA (DAPI, yellow). Scale bars, 5µm. **c.** Optical sections illustrating examples of end-on attachment, unattached, or lateral/merotelic attachment. Scale bar, 1 µm. **d.** The proportion of abnormal KT-MT attachments (unattached, lateral, and merotelic) in oocytes from YY, AA, and AY RCFs. 2-month-old (young) and 14-month-old (aged) wide type ICR mice were used. Box plots depict data distribution, where the mean and median are indicated by black square and center bar; upper and lower limits and whiskers are quartiles and 1.5 × interquartile range. One-way ANOVA, Tukey’s multiple comparisons test. \*\*\*\**P* < 0.0001; \*\*\**P* < 0.001; ns, *P* > 0.05.

**Extended Data Figure 12.**
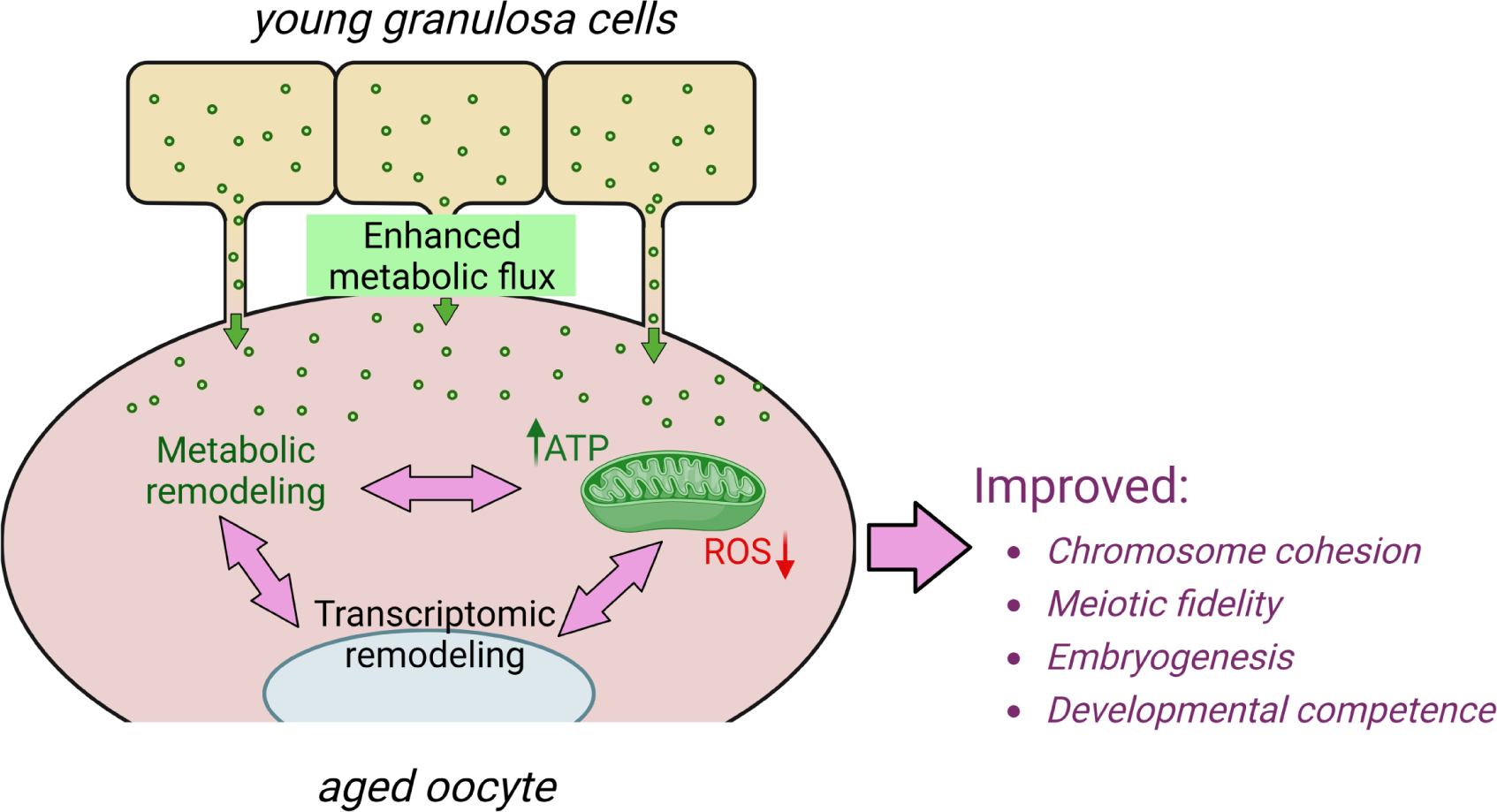
Scheme showing the proposed model of how young follicular somatic cells rejuvenate aged oocyte.

**Supplementary Video 1. Chimeric follicle generation process.** This video demonstrates a step-by-step example of creating a reconstituted chimeric follicle, highlighting the process of transplanting an oocyte into an r-follicle.

**Supplementary Video 2. An example of sister kinetochore pair distance measurement.** This video demonstrates the measurement of sister kinetochore pair distances in oocytes expressing 2mEGFP-CENP-C (green) and H2B-mCherry (red) to label kinetochores and chromosomes, respectively. See Methods for a detailed description of the measurement protocol.

**Supplementary Table 1.** Differential gene expression analysis *in vitro* oocytes versus *in vivo* oocytes.

**Supplementary Table 2.** Metabolomic profiling of oocytes from YY, AA, and AY RCFs,

**Supplementary Table 3.** Differential gene expression analysis in oocytes from AA RCFs versus YY RCFs.

**Supplementary Table 4.** Differential gene expression analysis in oocytes from AA RCFs versus AY RCFs.

**Supplementary Table 5.** Differential gene expression analysis in oocytes from AY RCFs versus AA RCFs.

